# A growth-survival trade-off quantitatively predicts microbial selection under periodic disinfection

**DOI:** 10.64898/2026.04.30.721825

**Authors:** Nerea Martínez-López, Adrián Pedreira, Míriam R. García, Frank Schreiber, Niclas Nordholt

## Abstract

Microbial populations frequently experience periodic lethal stresses from natural and anthropogenic sources, including routine disinfection in clinical, industrial, and domestic environments. However, periodic disinfection can rapidly select for tolerant strains with increased survival but reduced growth during permissive conditions, creating a trade-off that shapes competitive outcomes in microbial communities. Here, we develop a mathematical framework to quantify and predict selection between microbial strains competing for growth-limiting resources under periodic disinfection. The framework is validated through a competition experiment simulating periodic disinfection with benzalkonium chloride between a wild-type *Escherichia coli* strain and a tolerant mutant exhibiting increased survival but decreased growth. A minimal model incorporating growth rate during permissive conditions and disinfection survival quantitatively captures selection across different experimental scenarios, including uncertainty estimates propagated from parameter variance. We further provide analytical expressions and a web-based interface to determine selection outcomes and quantify the contributions of survival and growth rate to selection. Our framework establishes a quantitative basis for predicting when periodic disinfection shifts population composition towards tolerant strains and is generalizable to other lethal stresses, including antibiotic chemotherapy and physical inactivation, thereby contributing to our understanding of the impact of periodic selective pressures on microbial competition.

## 1 Introduction

Microbial populations in natural and built environments are frequently subjected to periodic lethal stresses that drastically reduce population size before growth resumes under permissive conditions. Some examples include antimicrobial chemotherapy, disinfection protocols, and other physical or chemical perturbations in engineered and natural systems. A common evolutionary response enabling microorganisms to evade lethal stress is tolerance, defined as the ability to survive transient exposure to otherwise lethal conditions without altering susceptibility to prolonged exposure, thereby distinguishing it from resistance [1–4].

Tolerance and resistance often incur fitness costs, typically manifested as reduced growth under permissive conditions, creating a trade-off between microbial survival under lethal stress and competitiveness in its absence [1, 5–8]. Understanding how growth-survival trade-offs influence microbial competitive outcomes and selection dynamics under periodic disturbances remains a central ecological challenge. Although prior studies have addressed different aspects of this question [9–14], there is still a need for experimentally testable and analytically tractable frameworks capable of quantitatively predicting microbial selection outcomes based on growth-survival trade-offs. Periodic disinfection provides a relevant and well-controlled system for developing and validating such frameworks, as it is commonly applied across clinical, industrial, and domestic settings to control microbial populations.

Disinfectants are biocides used to kill microorganisms across many settings, including hospital surfaces [15, 16], veterinary and animal husbandry [17–19], households [20, 21], food and feed processing [22–24], and water [25–27]. Their efficacy against target microorganisms is assessed using standardized test procedures [28] that require a specific reduction in viable cells following exposure of a microbial population. This criterion distinguishes disinfection from sterilization, which requires the practical eradication of viable cells [29]. Standard tests rely on surrogate microorganisms, enabling comparison between different products but disregarding variability among microbial strains and species occurring under real-world conditions [28].

Different strains within a microbial species can display distinct tolerance levels to disinfection [30, 31]. Furthermore, disinfectants are typically applied periodically, as in hygiene hospital protocols [15, 16, 32] or in surface disinfection for food processing [22, 23, 33]. However, experimental evolution studies have shown that spontaneously emerging tolerant *E. coli* mutants can be rapidly selected by periodic disinfection [34, 35]; such regimes could then foster natural selection of strains based on their tolerance if sterilization of the treated environment is not achieved. Periodic disinfection can thus increase the prevalence of tolerant strains in the exposed environment over time, potentially leading to disinfection failure and spread of disinfectant-tolerant strains across different settings [36–39].

One key prerequisite for natural selection of tolerance under periodic disinfection is a growth phase for those strains that preferentially survive biocidal exposure [6, 35, 40]. This may occur when nutrient supply is fluctuating, e.g., if surviving strains are transmitted from disinfected surfaces to nutrient-rich environments and then shed back onto surfaces, or if the disinfected setting is exposed to nutrients. Notably, tolerance to disinfectants and other antimicrobial substances often entails fitness costs under permissive conditions, such as reduced growth rate or prolonged lag [1, 2, 34]. These costs associated with altered antimicrobial susceptibility have been identified as important drivers for the selection of antibiotic resistance [41, 42].

From a practical perspective, it is desirable to set simple and measurable biological parameters to assess the outcome of disinfection protocols—e.g., in terms of log reductions or extinction thresholds [43,44]—and then use this information to predict the risk of selecting for resistant or tolerant strains within the treated setting to a level compromising disinfection efficacy. Mathematical modeling is a powerful tool for designing disinfection protocols, as it provides a structured framework for quantifying microbial pharmacodynamics [3,33,45–49] or competitive outcomes [12,14,33,34,45,46], and for identifying the key processes underlying the emergence and selection of resistance and tolerance. After validation, mathematical models can be used in practical applications to assess disinfection efficacy and explore optimal configurations to prevent selection of resistance or tolerance through *in silico* simulations or analytical expressions.

Fitness landscapes are increasingly used to study the emergence and fixation of resistant mutants under antibiotic exposure [47, 49–54], but these have several limitations when applied to periodic lethal stresses such as disinfection. Fitness landscapes are typically assumed to vary with antibiotic concentration according to the Mutant Selection Window (MSW), derived from the relationship between net growth rates and drug dose. Such models often lack analytical expressions for studying growth-survival trade-offs or rely on cell-level parameters that are difficult to measure, limiting their applicability in real-world scenarios. Another key limitation is the lack of a separation between periods of growth and lethal exposure. This separation is critical because strains with low tolerance but fast growth can outcompete tolerant strains during permissive periods [6, 34], so that the balance between these opposing selective pressures determines the population-level outcome. Although mathematical frameworks considering alternating phases of growth and lethal stress do exist [11,33,34,45,55,56], these do not provide predictions of selection outcomes from growth-survival trade-offs [33, 34, 45, 55, 56], lack analytical solutions [33, 34, 45, 55], or have not been validated experimentally [11, 33, 45, 55].

In this study, we develop a mathematical framework to assess the selection of microbial strains competing for common, growth-limiting resources under a periodic protocol, consisting of successive cycles alternating growth with disinfection periods. Our approach relies on a minimal model capturing the key mechanisms for microbial selection based on two easily measurable phenotypic traits: tolerance (i.e., disinfection survival), and fitness cost of increased tolerance, in terms of reduced growth rate in the absence of disinfectant (Figure 1). We provide analytical expressions to determine extinction and selection outcomes between the two strains, depending on the selective pressure exerted by competition and the strength of lethal stress (i.e., disinfectant dose), allowing us to dissect overall fitness into contributions from tolerance and growth. The model predictions are validated through competition experiments using two related *E. coli* strains obtained from a previous adaptive laboratory evolution experiment [34] that selected for tolerance to the disinfectant benzalkonium chloride: one susceptible ancestor and one tolerant mutant with a known fitness cost. To facilitate the application of the proposed framework in practical settings, we provide an accessible web-based interface, available at microracle.shinyapps.io/Microracle/?tab=PEplane.

**Figure 1:**
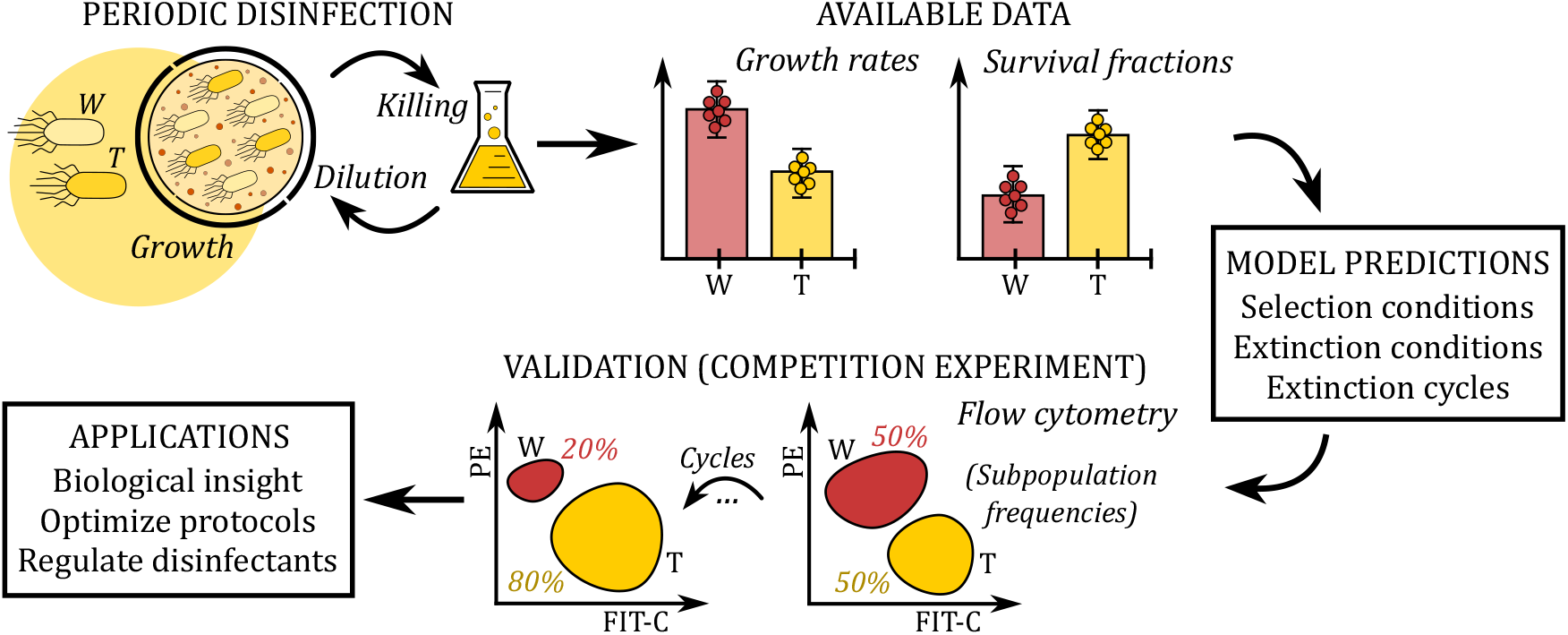
Understanding selection dynamics under periodic disinfection through mathematical modeling and experimental validation. Two microbial strains, W (red) and T (yellow), compete under periodic disinfection, alternating periods of growth and disinfectant exposure in each cycle. The growth rates determine the ability of the strains to compete for resources during growth, and cell survival after disinfection determines their ability to propagate to the next cycle. We assume that increased survival of T as compared to W comes at the expense of reduced growth. We developed a mathematical model to predict the selection dynamics of two such strains under periodic disinfection. The model was parametrized using growth and survival traits of strains isolated from a previous laboratory evolution experiment. The model predictions were validated with a competition experiment in which the strains were fluorescently tagged and monitored by flow cytometry over several cycles of growth and disinfection. The modeling approach has applications for understanding the selection dynamics of resistance or tolerance in microbial communities, for optimizing disinfection protocols to mitigate efficacy loss driven by the selection of resistant or tolerant strains, and for regulating disinfectants.

## 2 Materials and methods

### 2.1 Bacteria and culture conditions

All strains used in this study are derived from *E. coli* K12 MG1655, referred to as wild-type (W) throughout the text. The tolerant strain (T) was obtained from a previously published laboratory evolution experiment [34], designated there as S4. Both strains W and T were genetically engineered to express chromosomally encoded fluorescent proteins mCherry or YFP in a previous study [4]. Bacteria were cultured at 37 °C and shaking at 220 rpm in M9 medium. M9 medium was composed of 42 mM Na_2_HPO_4_, 22 mM KH_2_PO_4_, 8.5 mM NaCl, 11.3 mM (NH_4_)_2_SO_4_, 1 mM MgSO_4_, 0.1 mM CaCl_2_, 0.2 mM Uracil, 1 µg*/*mL thiamine, trace elements (25 µM FeCl_3_, 4.9 µM ZnCl_2_, 2.1 µM CoCl_2_, 2 µM Na_2_MoO_4_, 1.7 µM CaCl_2_, 2.5 µM CuCl_2_, 2 µM H_3_BO_3_) and 5 mM glucose. Unless stated otherwise, bacteria were cultured and treated in 1.5 mL 96-deep-well plates filled with 600 µL M9 per well.

### 2.2 Time-kill assays

To determine suitable conditions for the competition experiment, time-kill assays were performed. Strains W-YFP, W-mCherry, T-YFP, and T-mCherry were revived from freezer stocks in glass tubes filled with 2 mL M9 and cultured overnight. The cultures were diluted to a density of 1*×* 10^5^CFU*/*mL in M9 and grown for 24 h. Time-kill curves were recorded after BAC addition (Sigma-Aldrich, order no. 12060) to different concentrations (30, 40, 50, 60, 75 and 150 µg*/*mL). Cell counts were obtained by serial dilution in 1:10 steps in phosphate-buffered saline followed by drop plating of 7-10 µL on LB agar plates. The sampling times for measuring cell counts were 0, 5, 10 and 20 min. The results of the time-kill assay are shown in the Supplementary Information (Section S1). The conditions chosen for the competition experiments were 0, 30, 40, 50 µg*/*mL BAC and 10 min of exposure.

### 2.3 Competition experiment

The competition experiment was conducted in 96-well format, using a *Platemaster P220* (Gilson) 96-well pipettor to facilitate handling. Pre-cultures of W-YFP, W-mCherry, T-YFP, and T-mCherry were cultivated overnight and mixed at the desired ratios (1:1, 1:10^2^, 1:10^4^) to a final cell density of approximately 10^6^ CFU*/*mL in M9. For each mixing ratio and BAC concentration, 600 µL of this suspension was dispensed in triplicate into 1.5 mL deep-well plates. The deep-well plate was sealed with a cover plate and parafilm to prevent evaporation. Cells were then cultured for 24 h to reach the stationary phase. The following steps were repeated each day for the duration of the competition experiment: after cultivation, 10 µL from each well were withdrawn and diluted in 0.2 µm phosphate-buffered saline (PBS), followed flow cytometry to determine cell counts prior to killing (see Supplementary Information, Section S2). To initiate killing, 10 µL BAC solution were added to all wells to the desired final concentration (0, 30, 40, 50 µg*/*mL) and the plate was incubated at 37 °C and shaking at 220 rpm. After 10 min, 6 µL of the cell suspension were added to a new 96-deep-well plate containing 594 µL M9 per well, which was then sealed and incubated for 24 h as described.

### 2.4 Acquisition and treatment of flow cytometry data

The subpopulation frequencies of strains W-YFP, W-mCherry, T-YFP and T-mCherry were determined with flow cytometry before each disinfection step during the competition experiment. Flow cytometry was performed on the 96-well plate using a *CytoFlex S* flow cytometer (Beckman-Coulter) with a 96-well autosampler. Appropriate dilutions (typically 1:100) of the overnight cultures in filtered PBS were prepared in a 96-well plate, and 5 µL from each well was measured in the flow cytometer at a flow rate of 10 µL*/*min. This resulted in approximately 10^4^ *™*10^5^ events per well. Fluorescence of YFP was detected in the FITC-channel (excitation: 488 nm, emission: 525*/*40 nm). Fluorescence of mCherry was detected in the PE-channel (excitation: 561 nm, emission: 585*/*42 nm).

Flow cytometry data were automatically processed using MATLAB custom scripts implementing the following steps. First, undesired events (e.g., debris) were cleaned from the dataset by identifying outliers from FSC-A vs. SSC-A. Doublets and aggregates were also removed using SSC-H vs. SSC-A. Then, the signal intensities for YFP and mCherry in the FITC and PE channels were log-transformed, and a Gaussian mixture model was trained on the log-transformed datasets from wells on the first day, assuming three clusters: YFP, mCherry, and background. Finally, the cleaned datasets were classified into one of the three trained clusters, and the frequencies of YFP and mCherry were determined as the ratios of events in each cluster to the total number of events in the well.

To simplify the analysis and avoid duplicating results, flow cytometry data for the competition cases W-YFP vs. T-mCherry and W-mCherry vs. T-YFP were combined; the independence of strains’ survival under BAC exposure was confirmed by a two-way repeated-measures ANOVA (see Supplementary Information, Section S2).

### 2.5 The mathematical model

#### 2.5.1 Model formulation

We propose a minimal mathematical model to explain selection dynamics between two microbial strains, wild-type W and tolerant T, competing under periodic disinfection. The model reads (for detailed derivations and the model considering an arbitrary number of strains, see [57]):

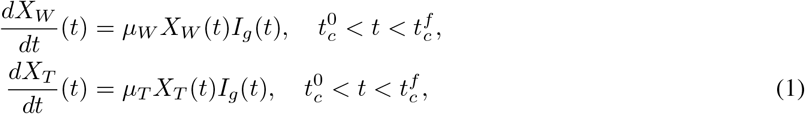

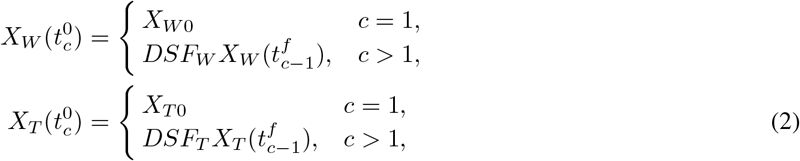

where subscripts *W* and *T* identify the strains throughout the model. *X*_*W*_ and *X*_*T*_ are the cell densities of the strains, which differ in their growth rates under permissive conditions, *µ*_*W*_ and *µ*_*T*_, and survival fractions after disinfection, *SF*_*W*_ and *SF*_*T*_. Each treatment cycle (*c*)—starting at 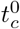 and concluding at 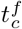 —consists of a growth period under competition for common, growth-limiting resources, followed by disinfection, and dilution by a factor *D* into fresh medium to start the next cycle.

During growth periods (Equation (1)), the strains grow exponentially until the next disinfection period starts or the total population reaches the medium’s carrying capacity, i.e., once resources are exhausted and cells enter the stationary phase (*I*_*g*_ = 1 during exponential growth, and *I*_*g*_ = 0 otherwise). When growth periods conclude, strains’ survival after disinfection—captured by the survival fractions (*SF*_*W*_ and *SF*_*T*_), implicitly depending on disinfectant dose and exposure time—and dilution into fresh medium set the initial cell densities for the next cycle (Equation (2)). Since disinfection is modeled as an instantaneous step, the *c*-cycle extends from 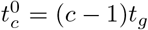 to 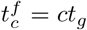, where *t*_*g*_ is the duration of the growth periods. At the first cycle, the initial cell densities are the strains’ inocula, *X*_*W* 0_ and *X*_*T* 0_, and the initial mixing ratio between the strains (initial ratio of T to W cells) is *T*_0_:*W*_0_ = *X*_*T* 0_*/X*_*W* 0_.

To mimic periodic disinfectant application, e.g., a daily disinfection protocol, the duration of each cycle is kept constant. However, the time during which cells actually grow is variable and depends on multiple factors. This can be calculated as:

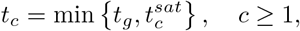

where 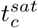 is the saturation time, i.e., the time necessary for the microbial population to exhaust the growth-limiting resource at the *c*-cycle. Consequently, the function *I*_*g*_ indicating the entry into the stationary growth phase in Equation (1) is:

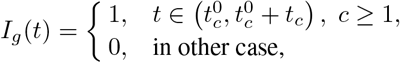

where the saturation time verifies:

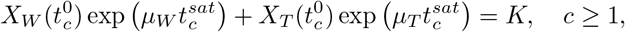

and *K* denotes the carrying capacity. Though this equation can only be solved numerically, we do not require any closed-form expression for the saturation time in most model derivations. This can be approximated as [57]:

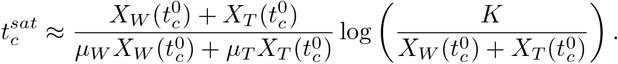

#### 2.5.2 Selection conditions

The model is used to predict selection outcomes between the two competing strains, W and T, under periodic disinfection. During permissive conditions, the ability of the strains to compete for growth-limiting resources relies on growth rate. The fitness cost of the tolerant T leads, therefore, to a selective disadvantage compared to the wild-type W under permissive conditions, since W grows faster and can consume the available resources before T becomes competitive enough. Here, we define the fitness cost (FC) of the tolerant T in terms of growth rate, as [47, 58]:

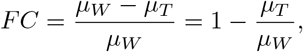

i.e., the fitness cost is the reduction in the growth rate of strain T (*µ*_*T*_) relative to the growth rate of W (*µ*_*W*_).

During disinfection, selective ability is determined by the survival fractions, so that the increase in tolerance confers a selective advantage to strain T over W. We quantify this survival advantage (SA) of the tolerant T as the log-ratio difference in the survival fractions between the two strains after disinfection (*SF*_*W*_ and *SF*_*T*_) and dilution by the factor *D*, relative to the wild-type W. That is:

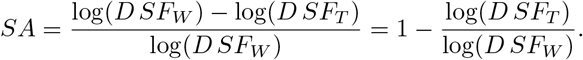

The trade-off between fitness cost and survival advantage of the tolerant T entirely determines the competitive outcomes between the two strains under periodic disinfection, leading to the following selection conditions [57]. Selection conditions (SC):

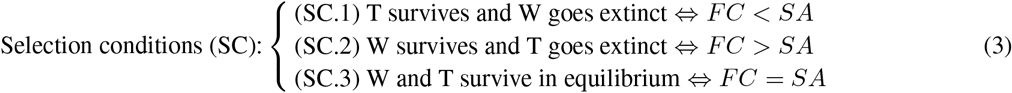

#### 2.5.3 Extinction conditions

The selection conditions determine which strain has the selective advantage under periodic disinfection; however, one strain could be selected over the other and still go extinct, which would mean that both strains are eventually eliminated. Then, the selection conditions cannot detect whether periodic disinfection successfully eliminates the microbial population.

Whether strains W and T persist or go extinct under periodic disinfection depends on how fast they can recover between disinfection steps. This is quantified by the equilibrium times:

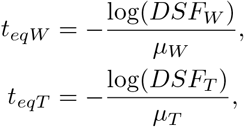

defined as the time required for the strains growing exponentially to replace the cells lost after disinfection and dilution. When the equilibrium time of a strain matches the time duration of the exponential growth phase at the *c*-cycle (*t*_*c*_), the cell density of the strain neither increases nor decreases over the cycle, i.e., net growth is zero.

Extinction under periodic disinfection occurs either because both strains go extinct in isolation—i.e., independently of competition—or because one or both strains surviving in isolation are driven to extinction by competition within a single treatment cycle. Assuming that a strain becomes extinct when its cell density falls below an extinction limit *X*_*e*_ (e.g., *X*_*e*_ = 1 CFU*/V*, where *V* is the culture volume), it goes extinct in isolation if: (i) the equilibrium time exceeds the growth time (*t*_*g*_), so that the strain cannot grow to compensate for cells removed after disinfection and dilution; or (ii) disinfection achieves a reduction in cell density from the carrying capacity to values below the extinction limit, in which case the strain is eliminated in a single treatment cycle. In contrast, extinction in competition occurs when a strain surviving in isolation cannot grow enough during the first cycle due to the presence of its competitor, so that it falls below the extinction limit after the first disinfection and dilution steps. This situation arises when competition effectively shortens the time available for recovery in the first cycle (*t*_1_). The possible extinction scenarios are summarized in the following extinction conditions [57].

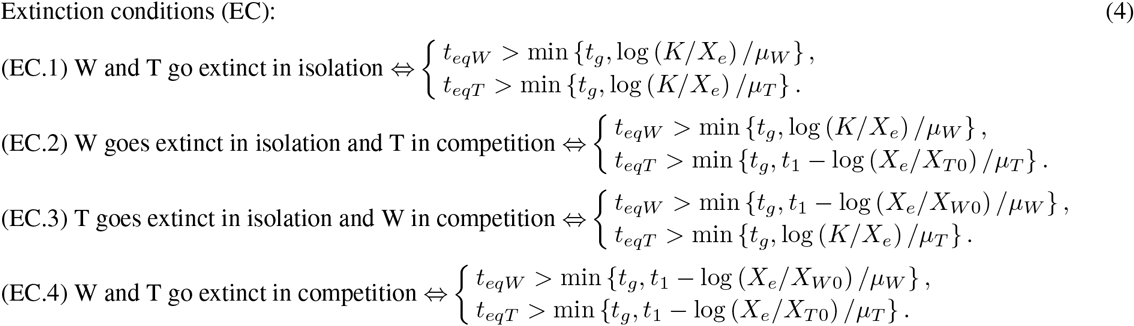

#### 2.5.4 Extinction cycles

If the extinction conditions predict the success of periodic disinfection, we can determine the number of cycles to eliminate the microbial population. The extinction cycle of the population is defined as the minimum number of cycles required to reduce the cell density of the two strains to values below the extinction limit (*X*_*e*_). This is [57]:

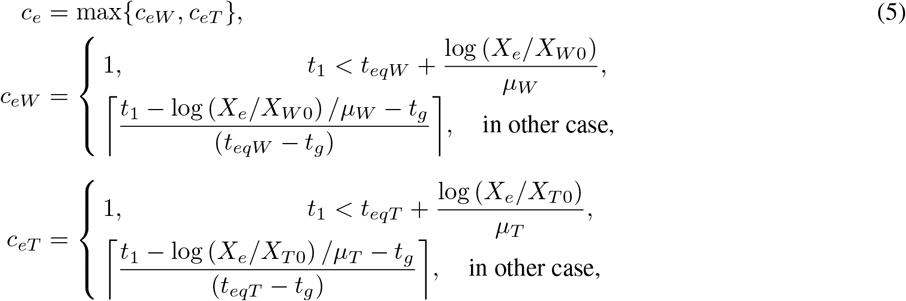

where⌈*x*⌉ denotes the ceiling function (the minimum integer greater than or equal to *x*).

When one strain survives periodic disinfection and outcompetes the other, we can approximate the extinction cycle of the strain going extinct. A closed-form expression is not available because we lack an analytical solution for the saturation times 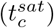, but we can approximate the extinction cycle reasonably by assuming that the surviving strain reaches the carrying capacity in the growth period immediately following the extinction cycle of its competitor. Then, if the tolerant T survives periodic disinfection and outcompetes the wild-type W, the extinction cycle of W can be approximated as [57]:

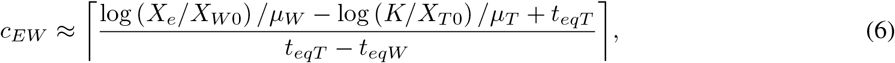

and, conversely, if strain W outcompetes T, the approximation for the extinction cycle of strain T is [57]:

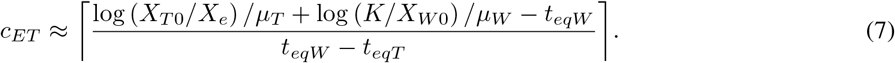

## 3 Results

### 3.1 A model-based approach for predicting selection under periodic disinfection

We combined mathematical modeling and experiments to quantify selection between a tolerant strain T and a wildtype strain W under periodic disinfection (Figure 2). Growth rates (*µ*_*W*_ and *µ*_*T*_) and survival fractions (*SF*_*W*_ and *SF*_*T*_) are determined experimentally for both strains in isolation and used to parametrize a minimal mathematical model (Figure 2A, Table 1). Experimental conditions, such as growth time, carrying capacity, and dilution factor, are required to set up the model. The mathematical model describes the dynamics of the cell densities for strains W and T under periodic disinfection at any cycle (Figure 2B). From the model, we derive analytical expressions to determine whether strains survive periodic disinfection, the conditions favoring selection of one strain over the other, or the disinfection cycle at which extinction occurs (Figure 2C; see Methods).

**Table 1:**
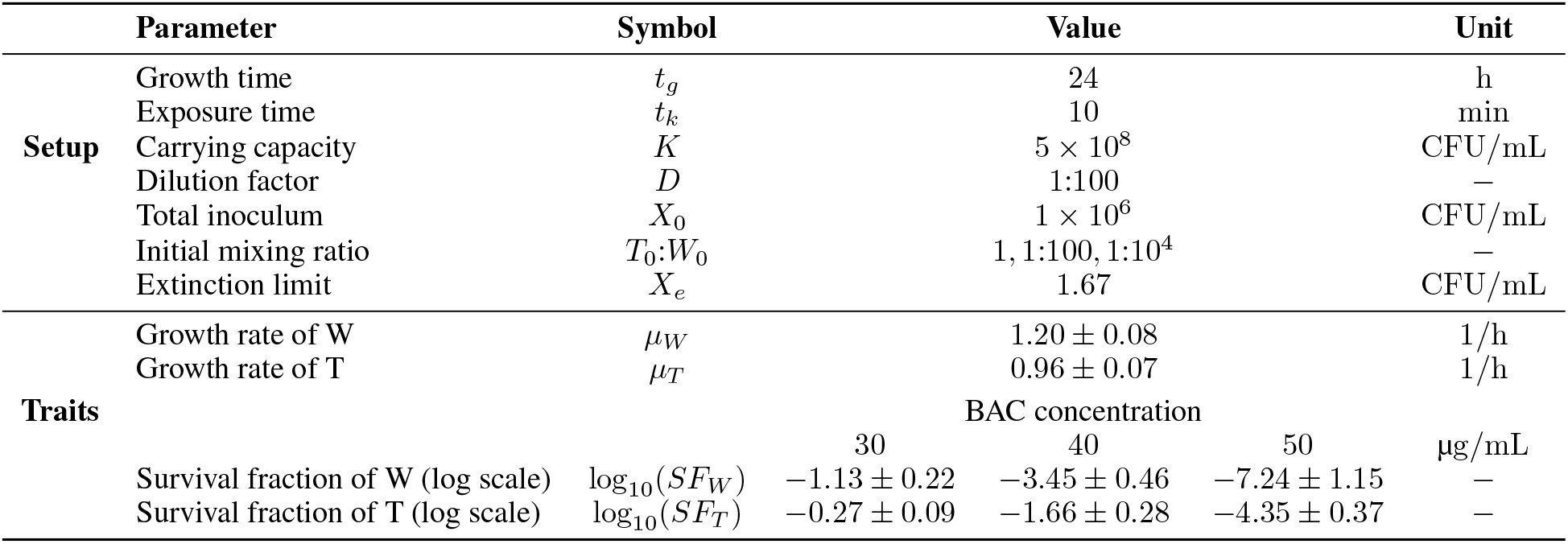
Parameters for simulating the competition experiment between the wild-type (W) and the tolerant (T) *E. coli* strains under periodic BAC disinfection, using the mathematical model in Equations (1)–(2). For details on parameter estimation from experimental data, see Supplementary Information (Section S3).

**Figure 2:**
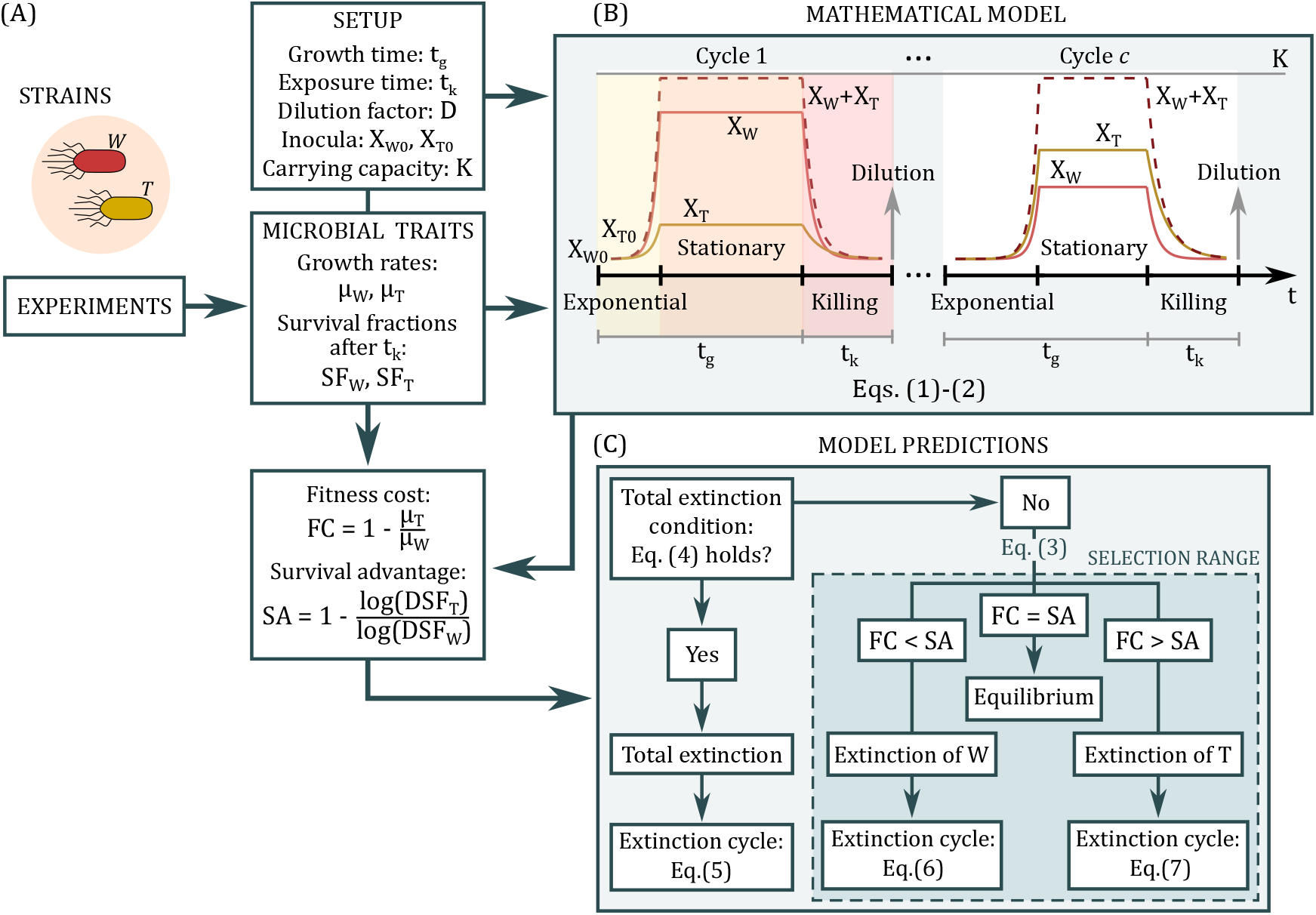
Approach for predicting selection under periodic disinfection using a mathematical model. (A) The microbial traits (growth rates and disinfection survival) of the competing strains, W and T, are determined from experimental data; (B) The microbial traits and setup parameters for periodic disinfection (e.g., exposure time or strains’ inocula) are used to parametrize the proposed mathematical model (Table 1); (C) The mathematical model (Equations (1)–(2)) can then be used to derive extinction conditions for determining disinfection success (Equation (4)). When disinfection is successful, we can estimate the extinction cycle of the mixed population (Equation (5)). Conversely, if bacteria evade periodic disinfection, two key parameters (fitness cost and survival advantage) determine which strain is selected (Equation (3)). Estimation of the key parameters relies on two main inputs: the microbial traits of the competing strains, and the setup parameters.

The mathematical model accounts for successive treatment cycles, consisting of a growth period under competition for common, growth-limiting resources, followed by disinfection and dilution into fresh medium to start the next cycle (Figure 2B). Growth proceeds until the next disinfection period starts or the total cell density reaches the medium’s carrying capacity as growth-limiting resources are exhausted. Hence, while the duration between disinfection periods remains constant, the time during which cells grow is variable and depends on the combined rate of resource consumption by the strains. Strain tolerance is quantified by the survival fraction, i.e., the fraction of cells (between 0 and 1)surviving after disinfection.

Because of its fast kinetics, disinfection is modeled here as an instantaneous step that reduces the cell density by the survival fraction, rather than a time-dependent kinetic. This has several advantages: (i) by considering only the disinfection outcome, the model is agnostic to the specific killing kinetics, which can be non-linear, (ii) the reduction in cell numbers at one time-point can be readily obtained experimentally, whereas the determination and interpretation of kill kinetics is more laborious, and (iii) regulatory requirements for disinfectants are defined by the reduction of viable cells, independent of mechanistic details.

### 3.2 The proposed modeling approach predicts selection across different scenarios

The growth rate of the tolerant T was approximately 20% lower than that of the wild-type W (0.96 1*/*h vs. 1.2 1*/*h; Figure 3A, Table 1). Killing increased with BAC concentration, and the survival fraction of T was higher than that of W at all the tested BAC concentrations (Figure 3A; Table 1). We then performed competition experiments simulating periodic disinfection between W and T, varying the BAC concentration to modulate the killing strength (Figure 3A). At the start of the competition experiments, the strains were mixed at different initial ratios (*T*_0_:*W*_0_) to investigate the effect of the initial abundance of the tolerant T on the selection outcome. Changes in the population composition were monitored by flow cytometry (Figure 3B; see Methods).

**Figure 3:**
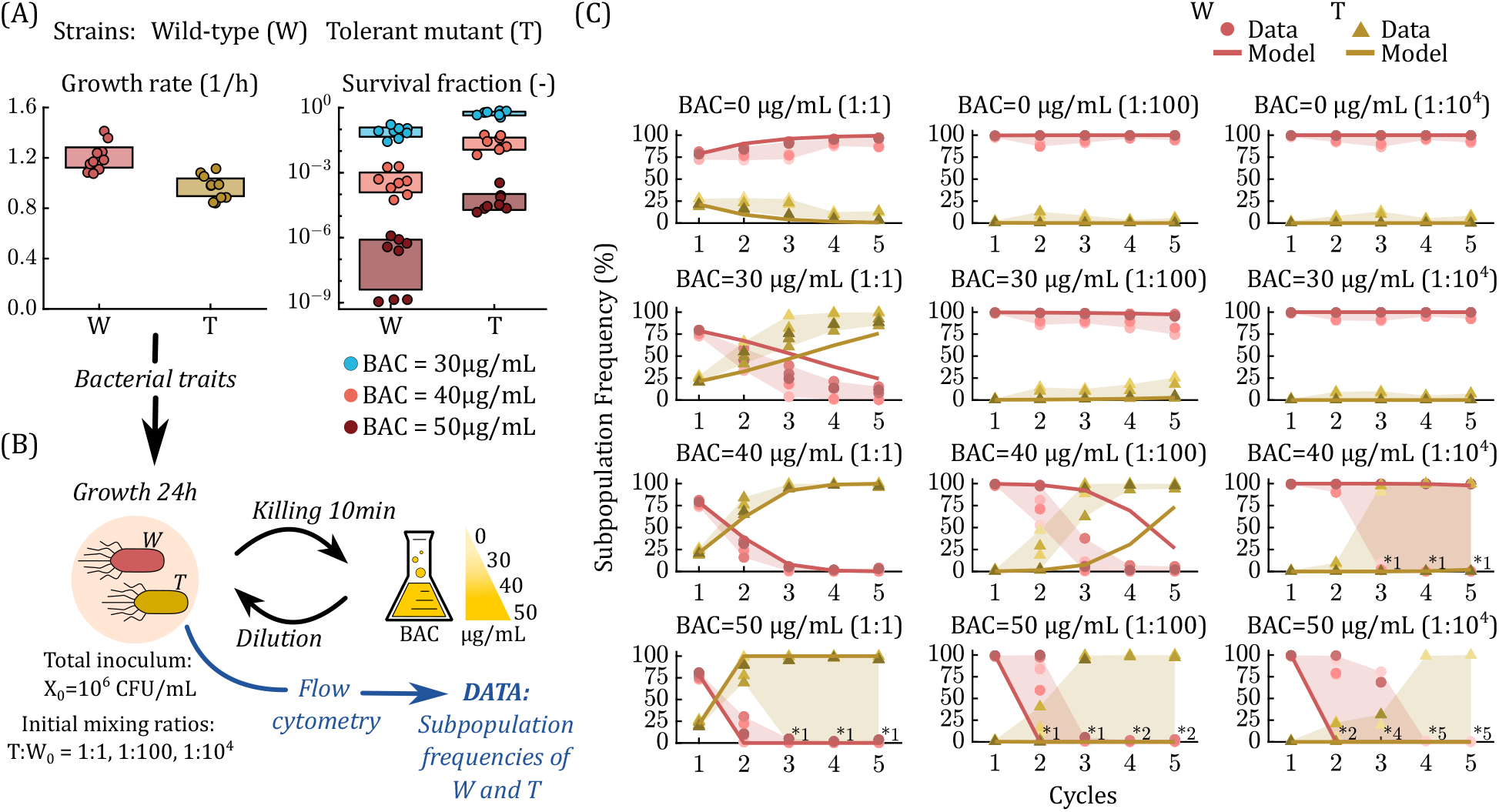
Predicting population dynamics under periodic disinfection across different scenarios. **(A) Microbial traits of the two competing strains.** Experimentally determined growth rate (*n* = 10 biological replicates) and tolerance after 10 min of BAC exposure (*n* = 8 biological replicates) for wild-type W and tolerant T (symbols), together with their 95 % confidence bounds (shading). Growth rates are taken from [34]; **(B) Setup of the competition experiment during periodic disinfection**. Overnight cultures of fluorescently labelled strains W and T were mixed at different ratios (*T*_0_:*W*_0_ = 1:1, 1:100, 1:10^4^). The mixed cultures were grown for *t*_*g*_ = 24 h, and then treated with different concentrations of benzalkonium chloride (BAC) for an exposure time *t*_*k*_ = 10 min. After disinfection, each cycle concludes by diluting the mixed cultures into fresh medium to start a new growth round. The frequency of the strains within the mixed cultures was determined at the end of the growth periods using flow cytometry; **(C) Comparison between experimental data and model predictions**. The subpopulation frequencies from model simulation (Equations (1)–(2); Table 1) are shown as continuous lines for the distinct combinations of initial mixing ratios (given in parentheses) and BAC doses. Symbols represent the biological replicates (*n* = 6) for the subpopulation frequencies detected experimentally at the end of each growth cycle (before disinfection), and asterisks represent population extinction, with the number of extinct replicates. Shading gives the variability range between experimental replicates.

In the absence of periodic disinfection (0 µg*/*mL BAC) the tolerant T is rapidly outcompeted by the wild-type W due to the fitness cost incurred by its increased tolerance (Figure 3C), even when the strains start from the same inoculum (*T*_0_:*W*_0_ = 1:1). As the BAC concentration increases, the survival advantage of T becomes increasingly determinant for selection. Conversely, when the initial mixing ratio decreases, the time until fixation of T is prolonged. At 50 µg*/*mL BAC, W is rapidly eliminated independently of T, due solely to killing and dilution. Thus, the importance of the selection component relying on cell survival can be adjusted through BAC concentration. Low BAC doses reduce the selective pressure on tolerance, thereby favoring the selection of the wild-type W, with the fastest growth. Conversely, high BAC doses favor the selection of the tolerant T, with slower growth but increased tolerance. The survival advantage of T compensates for its fitness cost, even at the lowest non-zero BAC dose (30 µg*/*mL). Only at the smallest inoculum of strain T (*T*_0_:*W*_0_ = 1:10^4^) the experiment’s outcome is not yet determined at 30 µg*/*mL BAC after 5 treatment cycles.

Model predictions become less accurate when the strains evolve close to extinction, because stochastic effects become significant and higher experimental variability is observed, i.e., at high BAC doses and low initial frequencies of T (Figure 3C). At the highest BAC dose, extinction of both W and T—i.e., total extinction—is detected in more replicates as the initial mixing ratio decreases. Total population extinction occurred in five of six experiment replicates at *T*_0_:*W*_0_ = 1:10^4^, indicating that W cannot survive periodic disinfection, independently of the presence of T. The population thus survives at the highest BAC dose only when there are enough T cells initially present in the mixed culture. Hence, the initial mixing ratio shapes the competitive ability of strain T at the start of the treatment, when it grows in competition with W. Specifically, competition with strain W during growth can prevent T from reaching a sufficient density before resources are exhausted and it will become extinct after disinfection and dilution. Taken together, the tolerant T has a selective advantage over the wild-type W under periodic disinfection with different killing strengths, and the mathematical model (Equations (1)–(2)) is able to satisfactorily capture the population dynamics during the competition experiment across different scenarios.

### 3.3 Selection as a trade-off between fitness cost and tolerance

We now use the mathematical model to derive conditions under which wild-type W and tolerant T are selected by periodic disinfection. These selection conditions can be computed directly from the competitive ability during growth and cell survival after disinfection, taking into account the commonly observed trade-off between fitness cost and tolerance under periodic lethal stress [1, 4, 34], which makes selection outcomes difficult to predict from individual traits alone.

We quantified this balance between growth and tolerance using two key parameters: fitness cost and survival advantage (see Methods). Briefly, the fitness cost (FC) captures the relative reduction in growth rate of strain T compared to W under permissive conditions, whereas the survival advantage (SA) captures the relative improvement in post-disinfection survival, accounting for killing and the subsequent dilution step. The trade-off between fitness cost and survival advantage defines the net selective advantage of T across a complete growth-disinfection cycle, which can be expressed as an overall fitness contrast (F = SA − FC), increasing with SA and decreasing with FC.

The selection conditions derived from the model (Equation (3)) yield three qualitatively distinct selection regimes [57]. First, when the survival advantage of the tolerant T exceeds its fitness cost (SA *>* FC), T is selected and W is driven to extinction; the increase in disinfection survival of T compensates for slower growth during resource competition, so its net increase per cycle exceeds that of W. This advantage can strengthen over successive cycles because increasing abundance and survival of strain T shorten the effective growth periods, thereby reducing the contribution of growth rate to total selection and shifting the balance further towards disinfection survival. Second, when the fitness cost dominates (SA *<* FC), W is selected and T goes extinct, as faster growth between disinfection periods outweighs any benefit from increased survival. Last, when fitness cost and survival advantage balance (SA = FC), neither strain has a selective advantage, and the system approaches coexistence at an equilibrium frequency.

These selection conditions are translated into a selection plane for the competition experiment where the fitness cost of T is represented against its survival advantage (Figure 4A). Mapping the experimentally determined parameters (FC and SA, Table 1) onto the selection plane visualizes the prediction that the tolerant T outcompetes the wild-type W for all tested BAC doses. At 30 µg*/*mL BAC, the strains are close to the selective equilibrium (total fitness F = SA - FC *≈* 0.07), which explains the prolonged competition (Figure 3C).

**Figure 4:**
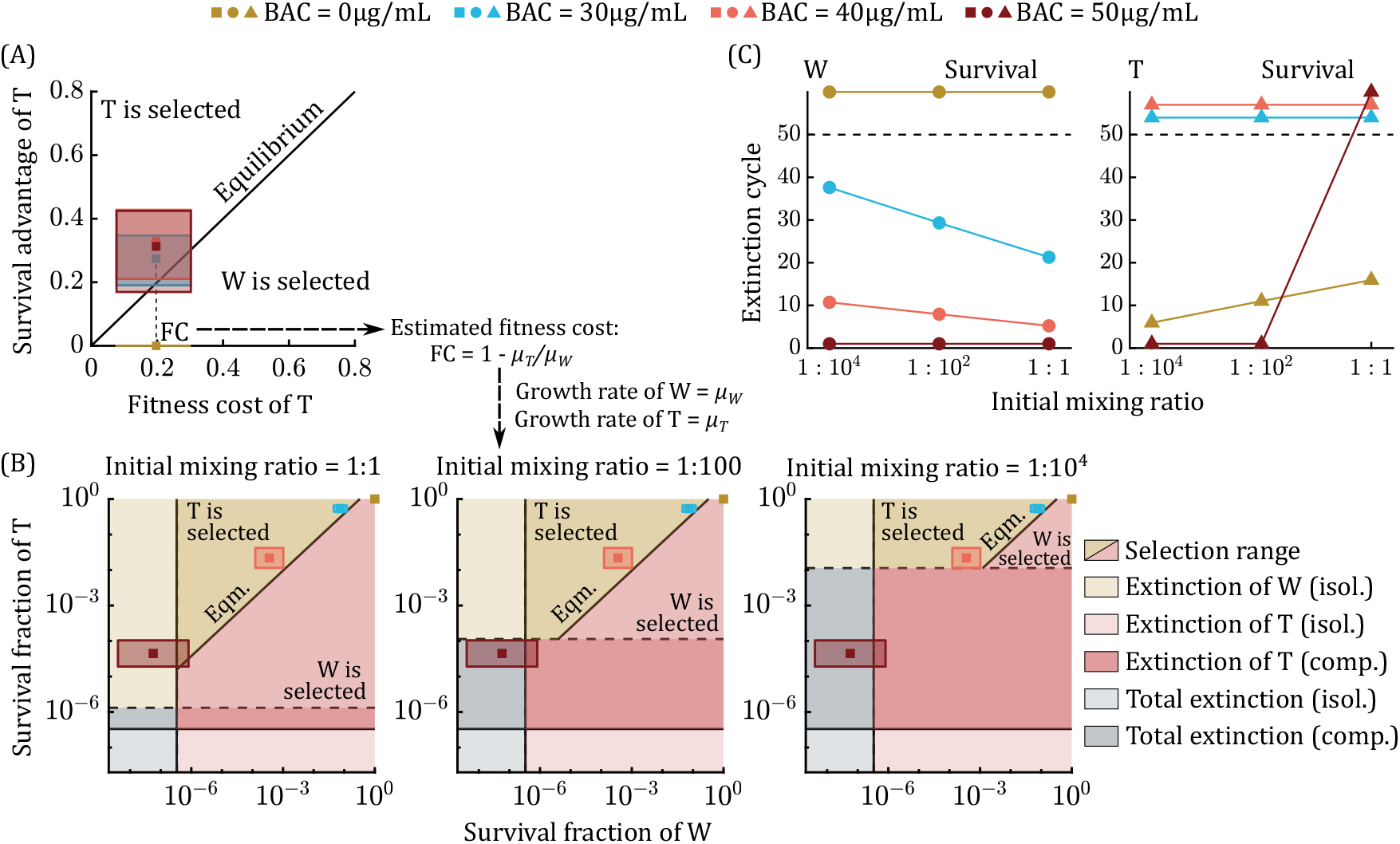
Model predictions of selection outcomes for wild-type W and tolerant T under periodic disinfection obtained from the experimental parameters. **(A) Selection plane for competition.** The fitness cost of the tolerant T is represented against its survival advantage (Equation (3)). Squares show the fitness cost and survival advantage at the different BAC doses obtained from the experimentally determined microbial traits (growth rates and survival fractions,Table 1). Shading represents the 95% confidence intervals derived from experimental parameters. **(B) Extinction planes at fixed fitness cost**. The survival fraction of wild-type W is represented against the survival fraction of the tolerant T when the growth rates are set to the experimentally determined values. Continuous lines limit the extinction ranges of the strains in isolation, and dashed lines define the bounds for extinction in competition. The extinction bound in competition for strains W largely coincides with the bound in isolation because the wild-type W has a large competitive advantage over the tolerant T during the first growth period and reaches a cell density close to that in isolation at the end of the period when the first disinfection step is performed. The range above the extinction bounds is the effective selection plane. The position predicted by the extinction conditions (Equation (4); Table 1) at the distinct BAC doses is represented by squares, and shading gives the uncertainty from the experimentally determined survival fractions; **(C) Predicted extinction cycles as function of disinfection strength and initial mixing ratio**. The extinction cycle predicted for strains W and T (Equations (6)-(7); Table 1) is represented against initial mixing ratio at the different BAC doses (dots for strains W and triangles for strain T). Extinction cycles above the dashed line indicate that the strains will not go extinct during periodic disinfection.

Together, these results show that periodic disinfection selects for tolerance only when the survival gain per cycle is sufficient to offset its growth penalty, with the boundary between competitive exclusion and coexistence defined by the equality between SA and FC.

From Equation (3), periodic disinfection selects for the tolerant T when the increase in the survival fraction relative to the wild-type W (*α* = *SF*_*T*_ */SF*_*W*_) verifies:

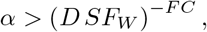

so that, in addition to intrinsic fitness costs, selection of T is negatively affected by dilution and BAC dose (which influences *SF*_*W*_). These parameters control the duration of the exponential growth phase within each treatment cycle, and thus modulate the contribution of growth rates to selection. Specifically, stronger dilution, i.e., decreasing the dilution factor (*D*), extends the exponential growth phase, thereby increasing the selective pressure exerted by growth-rate differences. Dilution thus favors the selection of W since it grows faster.

The minimum theoretical relative increase in survival fraction (*α*) at which the fitness of the tolerant T breaks even with the fitness of the wild-type W was calculated for typical values of fitness cost and dilution factor (Table 2). At a low fitness cost of T (5% reduction in growth rate relative to W), the survival increase necessary to outcompete W is also small (less than 5-fold) and approximately constant with changes in the dilution factor and BAC dose. When the fitness cost is 20%, the dependence of selection dynamics on the dilution factor and survival of strain W is much more evident. The minimum increase in the survival fraction of T ranges from *α ≈* 2.7 to *≈* 112, depending on the killing strength (i.e., BAC concentration) and the dilution factor. In all cases, the experimentally determined values for *α* were above the theoretical minimum (Figure 3A, Table 2). Hence, even at 30 µg*/*mL BAC, the model predicts that the tolerant T will eventually take over the population in our competition experiment (Figure 3C). For 50 µg*/*mL BAC, the experimentally determined increase in the survival fraction of strain T relative to W of 783 is well above the theoretical limit of selection of 70.6 for T (Table 2).

**Table 2:** Minimum relative increase in the survival fraction of the tolerant T (*α*) required to outcompete the wildtype *W*, depending on different fitness costs, dilution factors, and survival fractions of strains W (see Table 1). The experimentally determined values of fitness cost and increase in survival fraction of strains T (Figure 3A) are shown in the bottom row.

| | | Increase in survival fraction of T ($\alpha$) | | | |
| --- | --- | --- | --- | --- | --- |
| | | Dilution factor ($D$) | 30 | BAC ($\mu\text{g/mL}$)<br>40 | 50 |
| Fitness cost (FC) | 0.05 (Low) | 1:10 | 1.28 | 1.67 | 2.58 |
|  |  | 1:100 | 1.43 | 1.87 | 2.90 |
|  |  | 1:10<sup>3</sup> | 1.61 | 2.10 | 3.25 |
| | 0.2 (Intermediate) | 1:10 | 2.66 | 7.76 | $4.46 \times 10^1$ |
| | | 1:100 | 4.22 | $1.23 \times 10^1$ | $7.06 \times 10^1$ |
| | | 1:10<sup>3</sup> | 6.69 | $1.95 \times 10^1$ | $1.12 \times 10^2$ |
| | 0.5 (High) | 1:10 | $1.16 \times 10^1$ | $1.68 \times 10^2$ | $1.33 \times 10^4$ |
| | | 1:100 | $3.65 \times 10^1$ | $5.30 \times 10^2$ | $4.19 \times 10^4$ |
| | | 1:10<sup>3</sup> | $1.16 \times 10^2$ | $1.68 \times 10^3$ | $1.33 \times 10^5$ |
| FC Estimated | 0.197 | 1:100 | 7.19 | $6.16 \times 10^1$ | $7.83 \times 10^2$ |

The selection conditions do not apply when the entire population goes extinct under periodic disinfection; nevertheless, we derived conditions for determining total population extinction that depend on each strain’s traits rather than on overall fitness relative to the competing strain (Equation (4)). These conditions indicate that total extinction is still possible when neither of the strains goes extinct in isolation, because the final fraction of tolerant T cells, and therefore the effective population tolerance, is affected by resource competition [57]. Because many combinations of growth rate and survival fraction yield the same overall fitness, we fixed the growth rates to their experimentally determined mean values and constructed a reduced extinction plane representing the survival fraction of W against that of T (Figure 4B).

The predicted extinction plane captures the results of the competition experiments (Figure 4B), indicating that the tolerant T survives periodic disinfection and outcompetes the wild-type W for 30 and 40 µg*/*mL BAC. Additionally, the model predicts that W goes extinct in isolation at 50 µg*/*mL BAC, i.e., regardless of the competition with T. T is forced to extinction in competition with strain W as the initial mixing ratio decreases, even when W gets removed from competition due to extinction. As discussed before, the bound for extinction of the tolerant T in competition increases with the initial mixing ratio, so that the BAC dose to eliminate T decreases. This prediction provides an explanation for the extinction events at 50 µg*/*mL BAC during the competition experiments (Figure 3C). The confidence intervals for the position on the extinction plane overlap with the extinction bound (Figure 4B), suggesting that M is close to surviving periodic disinfection (see Supplementary Information; Section S6).

Predicting the number of cycles required to drive a resident wild-type strain to extinction during competition with a tolerant strain is relevant for designing periodic treatments or disinfection regimes. To this end, we derived an approximation for the extinction cycle of the wild-type W (Equation (6)). Again, we found good agreement between model predictions and experimental data (Figure 4C, Figure 3C). As experimentally observed, extinction of W occurs at earlier cycles when lethal stress becomes stronger, i.e. when BAC concentration increases. Furthermore, at fixed BAC concentrations, the time until extinction is shortened when the initial fraction of the tolerant T increases. In the cases where the outcome of the selection experiment was not fully determined after 5 cycles (Figure 3B at 30 µg*/*mL BAC), T is predicted to need between 20 and 30 cycles to drive W to extinction. In contrast, the number of cycles to drive T to extinction (Equation (7)) increases as the initial mixing ratio does. As observed in the competition experiment, strain W outcompetes T only at 0 µg*/*mL BAC, while the presence of W drives both strains to extinction after the first cycle at 50 µg*/*mL BAC, except when *T*_0_:*W*_0_ = 1:1.

## 4 Discussion

In this work, we developed a mathematical framework to quantify selection in microbial populations formed by a susceptible wild-type competing with a tolerant strain under periodic disinfection. The approach relies on a minimal dynamical model (Equations (1)–(2)) capturing the key mechanisms for microbial selection under successive growth and disinfection periods, based on: (i) growth rates, determining the competitive ability for common, growth-limiting resources during permissive conditions, and (ii) survival fractions after disinfection, quantifying disinfectant tolerance.

Our framework includes analytical conditions to determine selection outcomes (selection conditions, Equation (3)) or disinfection success (extinction conditions, Equation (4)), enabling direct quantification and prediction of the contributions of growth rate and tolerance towards selection under periodic disinfection. Both parameters can readily be obtained from strains isolated from the setting to be disinfected. The approach also allows us to determine the number of disinfection cycles to eliminate the microbial population when periodic disinfection is predicted to be effective (Equation (5)), or to approximate the number of cycles needed for competitive exclusion of one of the strains when disinfection is unsuccessful (Equations (6)–(7)). To facilitate the implementation of the proposed framework in practical applications, we developed a web-based tool (microracle.shinyapps.io/Microracle/?tab=PEplane) for analyzing selection and extinction outcomes.

The mathematical model predicted the experimentally observed selection dynamics across different scenarios for a competition experiment between a wild-type *E. coli* strain and a benzalkonium chloride (BAC)-tolerant mutant under periodic disinfection, based on independently measured growth rates and survival fractions (Figure 3). Our analysis showed that selection outcomes during periodic disinfection can be explained by the trade-off between fitness cost (reduced growth rate) and survival advantage (increased disinfection survival) of the tolerant *E. coli* strain [34] (Figure 4A). Strong selective pressures exerted by high disinfectant doses favor the selection of the tolerant strain, though the survival advantage needed to be selected over the wild-type increases with disinfectant dose (Table 2). This is because stronger killing leads to lower cell densities, which prolongs the next growth period and, in turn, amplifies the fitness costs during growth (Table 2). Similarly, the dilution factor modulates the relative importance of fitness costs [10, 59]. Due to these effects, higher disinfectant doses which do not achieve full eradication of a mixed population may prevent tolerant strains with reduced growth rate to become dominant in the population upon regrowth.

Interestingly, although the tolerant strain in our competition experiments would be able to survive treatment with the highest BAC concentration, the competition with the wild-type results in extinction of the entire population (Figure 4). This occurs because resource competition with the faster-growing wild-type prevents the tolerant strain from reaching a sufficient density to survive the subsequent disinfection step. As a consequence, total extinction can occur even when neither strain would go extinct in isolation, reducing the effective population tolerance by resource competition. This has a practical implication for disinfection design: competition with susceptible strains during growth can contribute to disinfection success, so that the presence of a fast-growing susceptible majority may be more effective at eliminating a tolerant subpopulation than increasing the disinfectant dose alone.

Our framework also predicts when adaptation becomes critical. For a given disinfection protocol targeting a specific log-reduction—as required by standard efficacy tests [28], for example—the derived selection conditions define the maximum increase in survival fraction a tolerant strain can acquire before it evades disinfection or outcompetes the susceptible population. This allows practitioners to define quantitative tolerance thresholds: if monitoring reveals that strains isolated from a disinfected environment exceed these thresholds, the disinfection protocol should be adjusted. Conversely, the framework could inform the design of standardized efficacy tests by incorporating selection-based criteria alongside the conventional requirement of a minimum log-reduction, thereby accounting for the risk that tolerant strains may stablish over repeated application. The proposed model of periodic disinfection is deterministic (identical parameter values produce identical dynamics) and therefore cannot account for stochastic dynamics arising when the microbial population is close to extinction [43, 60]. The most notable differences between model predictions and experimental data were thus observed in scenarios where the outcome of periodic disinfection is highly stochastic, with only some of the biological replicates going extinct (Figure 3C). Nevertheless, our modeling approach leverages the uncertainty of experimental parameters to account for uncertainty in model predictions. This allows the competition experiment to be mapped onto the selection and extinction planes predicted by the model, along with the expected uncertainty in these predictions (Figures 4A and 4B). Our framework can then be used to assess the efficacy of periodic disinfection in worst-case scenarios (e.g., when the fitness cost of tolerance is the lowest experimentally observed) when extinction and selection outcomes are highly uncertain. Because uncertainty in the experimentally determined traits to parametrize the model propagates into model predictions, minimizing parameter uncertainty improves prediction accuracy, e.g., through optimal experimental design [24, 61, 62].

The mathematical model proposed here can be extended to include other microbial traits that may influence selection outcomes under periodic disinfection. First, we assume exponential growth during growth periods until the population reaches the stationary phase due to exhaustion of the growth-limiting resources, so that strains’ competition is modeled indirectly [9, 10, 13]. The importance of growth periods in selection dynamics is thus modeled through the exponential growth rates of the isolated strains, which determine their ability to compete for resources and define the fitness cost of tolerance. Consequently, our model does not consider resource concentration dynamics (typically modeled using Monod kinetics [34, 46]), which enables analytical solutions, but disregards the influence of metabolic efficiency parameters, such as resource efficiency (yield) [9, 10, 13] or uptake affinity [63]. While some studies indicate that resource efficiency is not under direct selection [10, 12] and provide selection conditions ignoring this parameter [10], other works suggest that adjusting metabolic efficiency could be a relevant mechanism for the selection of resistant strains at sub-inhibitory drug concentrations [63, 64].

Another important parameter currently not included in our model is lag time, i.e., the time before growth resumes in fresh medium after starvation. Because lag can convey tolerance to lethal stress, adaptation of lag time could be a strategy adopted by tolerant subpopulations during favorable environmental conditions to anticipate future adverse stimuli [1, 7, 65]. Additionally, exposure to acute physical or chemical stresses can induce extended lag, thereby affecting competition dynamics [65, 66]. Since no growth occurs during lag, tolerance by lag is also an example for a trade-off between reproduction and tolerance: remaining in lag phase may increase survival, whereas fast growth resumption is an advantage during permissive conditions [1, 6, 9]. Interestingly, this trade-off can partly be alleviated through phenotypic heterogeneity in lag times [67].

Additionally, our model can be extended to systems where the lethal stress intensity changes over time—e.g., antibiotic decay within a patient or fluctuating disinfectant concentrations—by incorporating concentration-dependent killing [57, 68, 69]. Also, phenotypic heterogeneity is not considered, although it may affect selection dynamics. For instance, tolerance is implemented as a phenomenological measure of survival fraction after a fixed disinfection time. Microbial killing dynamics are often shaped by cell-to-cell heterogeneity in properties characterizing survival, which may underpin phenomena such as heteroresistance [42,60,70] or persistence [34,35,40]. Implementing killing dynamics would allow *in silico* simulations of disinfection protocols by changing disinfectant dose and exposure time [57]. However, this requires additional model parameters [35, 60, 71] that are not always experimentally accessible, especially in application conditions. While the focus of this study is on selection dynamics between competing strains already coexisting in a given environment, another future direction is the implementation of tolerant strains entering the population through *de novo* tolerance mutations.

Because the mathematical model is agnostic to the specific cause of killing and requires only two measurable parameters, our framework applies to any periodic lethal stress imposing a growth-survival trade-off on competing microbial strains. Beyond disinfection, relevant scenarios include antibiotic chemotherapy, where repeated dosing creates cyclic selection between tolerant and susceptible subpopulations; competitive microbial interactions mediated by inhibitory substances; physical perturbations such as heat or UV treatment in engineered systems; and freeze–thaw or desiccation–rewetting cycles in natural environments. In each case, the analytical expressions derived here can predict selection outcomes provided that estimates of strain-specific growth rates and survival fractions are known. Together, the proposed framework provides a general quantitative tool for understanding how periodic lethal stresses shape microbial competition through the interplay of survival and growth.

## 5 Conflicts of interest

The authors declare that they have no competing interests.

## 6 Funding

This work was supported by the Horizon Europe project “STOP: Surface Transfer of Pathogens” funded by the European Union’s Horizon Research and Innovation Program under Grant Agreement No. 101057961. This work was also funded by Spanish National Research Council (20213AT001, 02570E193 and iMOVE programme) and by MCIN/AEI/10.13039/501100011033 and “ERDF A way of making Europe” (PID2022-136817OB-I00).

## 7 Data availability

The data and code supporting the findings of this study are available at Nerea Martínez López, Adrián Pedreira, Míriam R. García, Frank Schreiber, & Niclas Nordholt. (2026). Experimental Datasets and Code for: “A growth survival trade-off quantitatively predicts microbial selection under periodic disinfection.” (v1.1). Zenodo. https://doi.org/10.5281/zenodo.19910150

The web-based interface implementing the calculations described in this study is available at: microracle.shinyapps.io/Microracle/?tab=PEplane.

## 8 Author contributions statement

Nerea Martínez-López (Conceptualization, Methodology, Softw-are, Validation, Formal analysis, Investigation, Data Curation, Visualization, Project Administration, Funding Acquisition), Adrián Pedreira (Software, Validation, Visualization), Míriam R. García (Conceptualization, Resources, Supervision, Project Administration, Funding Acquisition), Frank Schreiber (Concep-tualization, Resources, Supervision, Project Administration, Funding Acquisition), Niclas Nordholt (Conceptualization, Methodology, Validation, Investigation, Data Curation, Supervision, Project Administration). All authors contributed to writing the original draft and editing.

The authors confirm that AI tools did not generate content, influence the scientific reasoning, or contribute to the originality of the work. Large language models were used for reviewing spelling and grammar during the editing process.

## supplementary Information

### S1 Quantification of tolerance under disinfection with benzalkonium chloride

The survival of the wild-type *E. coli* strain (W) and the tolerant *E. coli* mutant (T) under benzalkonium chloride (BAC) disinfection was quantified by performing time-kill curves (TKCs) for the cells in the stationary phase (Figure S1A). The strains were previously shown to increase their survival (tolerance) when treated with BAC during the stationary growth phase compared to the exponentially growing cells, where strain T mutated from W to increase its tolerance level at the expense of a fitness cost (reduced growth rate) [1]. Cell survival was then quantified in the stationary phase to consider the tolerance of the strains and the survival advantage conferred by the mutation on strain T. Since the strains were labeled with protein YFP or mCherry to sort cells with flow cytometry during the competition experiment, the stationary TKCs were obtained for each combination of strain (W or T) and label (YFP or mCherry) separately to ensure that fluorescent labeling did not alter the results. The independence of the TKCs on the fluorescent tag was confirmed by a two-way repeated measures ANOVA, so that data were combined for the two labels and we considered a unique dataset per strain (Figure S1B).

The results of the stationary time-kill assay confirmed the increase in disinfection survival for the tolerant T compared to the wild-type W (Figure S1B). The survival advantage of strain T is only slightly appreciable at low BAC doses, but the differences in disinfection survival between the two strains become more significant as the concentration increases. For 50 µg*/*mL BAC, particularly, all the replicates for strain T survive when those of W fall below the low detection limit (LDL) at the end of the experiment (20 min). However, neither of the replicates for the tolerant T survive after disinfection for BAC concentrations higher or equal than 60 µg*/*mL. In general, BAC doses above 60 µg*/*mL decrease the cell density of the strains to values below the low detection limit in less than 5 min of BAC exposure (150 µg*/*mL BAC was also tested, but is not shown in Figure S1 as it overlaps with the curves at 75 µg*/*mL, thus providing no additional information). The survival advantage of the tolerant T is then restricted to a BAC concentration window, such that the differences in cell survival between strains W and T are maximum in the range 30 − 60 µg*/*mL BAC.

The survival increase of the tolerant T relative to the wild-type W after 10 min of disinfection (exposure time for the competition experiment) varies from approximately 10-fold at 30 µg*/*mL BAC, to more than 1000-fold at 50 µg*/*mL BAC (Figure S1C). The survival advantage of strain T over W thus depends on the BAC concentration, so that increasing the BAC dose also increases the differences in cell survival between the strains. The survival fraction decreases with the BAC concentration more drastically for the wild-type W than for the tolerant mutant T, which reduces its tolerance to the BAC pressure in a more gradual fashion.

### S2 ANOVA for independence of the time-kill curves on fluorescent tags

We performed a two-way ANOVA with repeated measures to test the hypothesis that the stationary time-kill curves (TKCs) for strains W and T depend on the fluorescent tag (YFP or mCherry). The two factors explaining the measured response (cell density) were strain and label, and the response was determined on each biological replicate (*n* = 4 per combination of strain and label) at multiple times and BAC doses (repeated measures). Interactions between the two factors were also considered to analyze possible conditional effects on the TKCs.

The results (Table S1) indicate a strong dependence of cell density on time and BAC dose, with associated *p*-values (1.5 × 10^−17^ and 2.6 × 10^−17^, respectively) fairly below the accepted limit (0.05). This is expected, as cell survival decreases with exposure time at a rate depending on the BAC dose. The measured response is also heavily influenced by the strain (associated *p*-value of 2.6 × 10^−6^), as the tolerant T shows increased survival compared to the wild-type W at all tested BAC doses and sampling times. On the contrary, the *p*-value for the dependence of the TKCs on the fluorescent tag (*p* = 0.96) is nearly one, providing strong evidence that cell survival under BAC exposure remains unaffected by staining.

Similarly, there is no evidence for conditional effects on the measured cell density resulting from interactions between the two factors (associated *p*-value of 0.51). Strain and label are thus weakly correlated, and the combined effect on the response can be satisfactorily explained through the independent effects of each factor (main effects); this means that the relationship between cell survival and strain remains unaffected by the label, and vice versa. Last, the strain factor is highly correlated with time and BAC dose (associated *p*-values of 1.7 × 10^−8^ and 4.4 × 10^−6^, respectively) indicating important interaction effects between strain, time and BAC concentration on the TKCs. The label factor, contrarily, does not show any correlation with time or BAC dose (*p*-values of 0.98 and 0.99, respectively).

**Figure S1:**
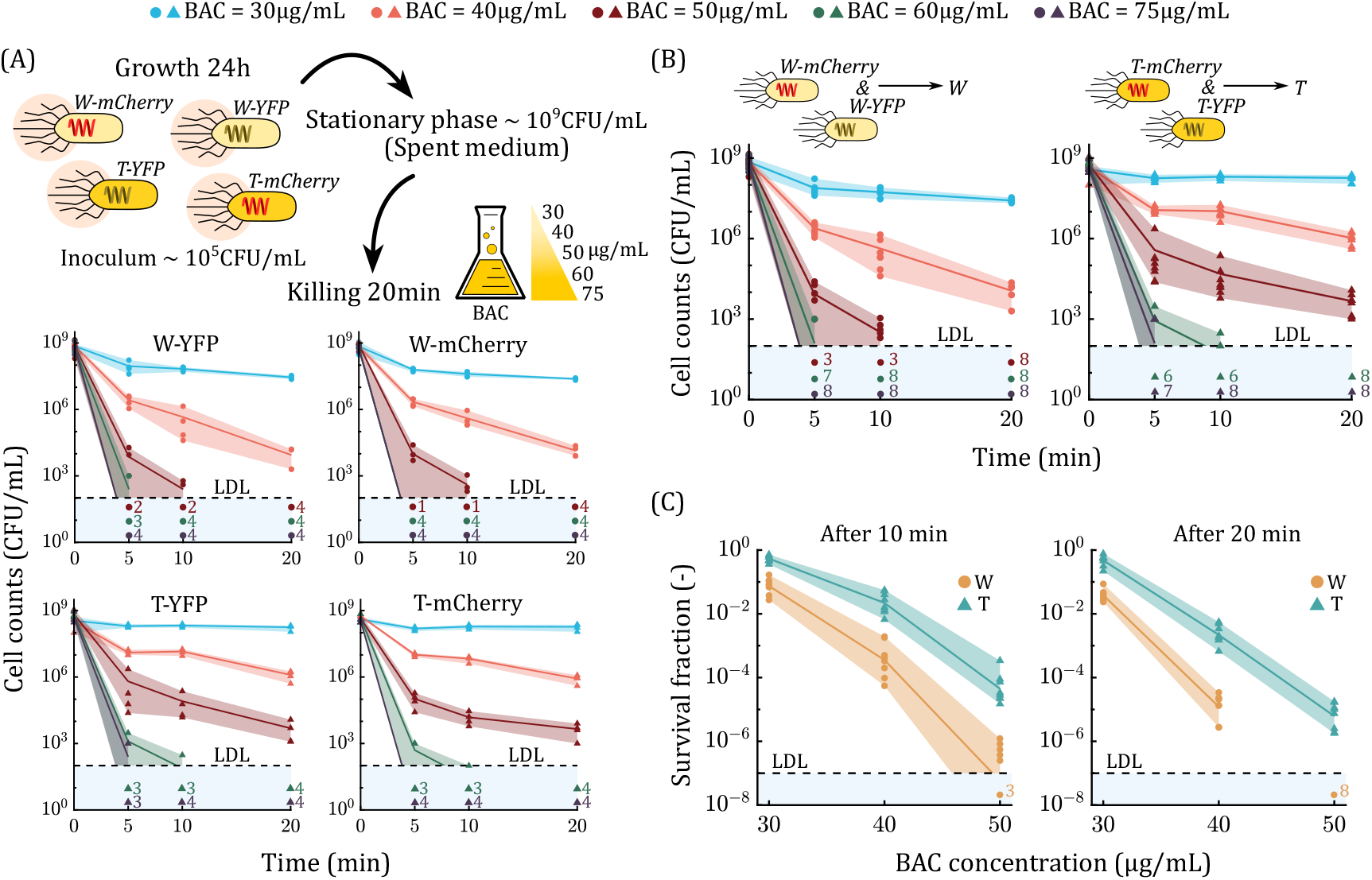
Results of the stationary time-kill assay. (A) Determination of the time-kill kinetics against BAC for wild-type (W) and tolerant (T) *E. coli* strains with fluorescent tags. Overnight cultures of the strains were labeled with YFP or mCherry and grown independently for 24 h (*n* = 4 biological replicates per combination of strain and label). Bacteria reproduced till the stationary phase, once the growth-limiting resources in the medium were exhausted (spent medium). The bacterial response against BAC disinfection was then tested at 30, 40, 50, 60 and 75 µg*/*mL for each combination separately. Cell counts measured at 0, 5, 10 and 20 min are represented by symbols (dots for strain W and triangles for strain T), shading represents the variability between replicates, and the continuous line is the mean. The numbers indicate the number of replicates for which counts below the low detection limit (LDL = 100 CFU*/*mL) were detected; **(B) Stationary TKCs for strains W and T**. The TKCs proved to be independent of fluorescent labeling by a two-way repeated measures ANOVA, so that data were combined for the two labels (*n* = 8 biological replicates per strain); **(C) Comparison of cell survival for strains W and T**. The survival fractions after 10 min (exposure time for the competition experiment) and 20 min of BAC disinfection are represented by symbols. Shading gives the variability between replicates, and the continuous line represents the mean.

In view of the results, we concluded that the fluorescent tag has a negligible impact on cell survival along the time-kill assay, independently of the strain. Therefore, wild-type W cells labeled with YFP or mCherry can be considered identical, and analogously for tolerant T cells.

### S3 Parameters for model simulation

The necessary parameters for simulating the competition experiment using the proposed mathematical model (see Methods in the main text) were chosen as follows.

Growth periods lasted for *t*_*g*_ = 24 h. The initial mixing ratio (*T*_0_:*W*_0_) was varied between 1:1 (equal competition, one T cell per W cell), 1:100 (slight initial advantage for strain W, one T cell per 100 W cells), and 1:10^4^ (high advantage for strain W, one T cell per 10^4^ W cells in the initial mixed culture). The total inoculum (initial density of W and T cells) was set to a relatively high value, *X*_0_ = 1 × 10^6^ CFU*/*mL, to avoid the removal of strain T by dilution when it starts from the lowest initial mixing ratio (1:10^4^). Resources were adjusted in the culture medium to obtain a total cell density of approximately 1 × 10^9^ CFU*/*mL in the stationary phase. Since no significant differences in cell density were observed between strains W and T at the start of the stationary time-kill assay (see Figure S1), we assumed the same resource efficiency (yield) for both strains, and worked with the carrying capacity as a setup parameter (note that the culture conditions for the time-kill assay and the competition experiment were identical, except that the inoculum was increased from 1 *×* 10^5^CFU*/*mL to 1 *×* 10^6^CFU*/*mL in the last case). The carrying capacity was thus calculated as the mean cell density at the start of the time-kill assay for the two strains, resulting in *K* = 5 *×* 10^8^ CFU*/*mL.

**Table S1:** Results (*p*-values) for the two-way repeated measures ANOVA to test independence of the time-kill curves on fluorescent tags. The two factors are strain (W or T) and label (YFP or mCherry), and interactions between the two factors (Strain:Label) are considered. The response variable (cell density) is measured on the same biological replicate (*n* = 4) at multiple sampling times and BAC concentrations (repeated measures).

|  |  | Factors |  |  | Repeated measures |  |
| --- | --- | --- | --- | --- | --- | --- |
| | | Strain | Label | Strain:Label | Time (min) | BAC ($\mu\text{g/mL}$) |
| **Response** | Cell density ($\log_{10}$ (CFU/mL)) | $2.6 \times 10^{-6}$ | 0.96 | 0.51 | $1.5 \times 10^{-17}$ | $2.6 \times 10^{-17}$ |
| **Factors** | Strain | | | | $1.7 \times 10^{-8}$ | $4.4 \times 10^{-6}$ |
|  | Label |  |  |  | 0.98 | 0.99 |
|  | Strain:Label |  |  |  | 0.40 | 0.49 |

The exposure time for disinfection periods was *t*_*k*_ = 10 min, and the tested BAC concentrations were 0, 30, 40 and 50 µg*/*mL. After each disinfection round, 6 µL of the mixed culture was diluted into 594 µL of fresh M9 medium, obtaining a dilution factor of *D* = 1:100. Last, we considered a limit *X*_*e*_ on the cell density below which the strains are assumed to be extinct. This limit was set to 1 CFU in a culture volume of *V* = 600 µL, so that *X*_*e*_ = 1.67 CFU*/*mL.

On the other hand, the phenotypic traits of strains W and T used for model-based simulation of the competition experiment were determined as follows. The exponential growth rates (*µ*_*W*_ and *µ*_*T*_) were calculated as the mean of *n* = 10 biological replicates for a growth assay performed in [1] under the same culture conditions considered here. Confidence intervals for the growth rates were calculated directly from the sample by assuming normally (Gaussian) distributed measurements of the growth rates around the mean. The survival fractions of the two strains (*SF*_*W*_ and *SF*_*T*_) after 10 min of BAC disinfection were obtained from the preliminary time-kill assay (see Section S1, Figure S1). The estimated survival fractions (in log-scale) were then calculated as the mean of *n* = 8 biological replicates for the time-kill assay, and the confidence bounds were obtained by assuming a Gaussian sample in the log-scale. Cell counts below the low detection limit (LDL = 100 CFU*/*mL) were also considered for estimating survival fractions, as these contain additional information for parameter estimation [2], and treated as zero counts. The survival fractions estimated with this procedure were compared with those estimated by the left-censoring approach and by assigning values equal to LDL or 0.5LDL to cell counts below the low detection limit, yielding fairly similar results in all cases.

Note that the stationary time-kill assay was performed separately for strains W-YFP, W-mCherry, T-YFP, and T-mCherry (*n* = 4 biological replicates each) to ensure that fluorescent labeling did not alter disinfection survival. This was confirmed by a two-way repeated-measures ANOVA (Section S2, Table S1), so the time-kill data of strains W and T were combined for the two fluorescent labels (resulting in *n* = 8 biological replicates per strain). Additionally, the dependence of growth rate and survival fraction on the fluorescent label was tested by fitting the mathematical model to the competition experiment data for the competition cases W-YFP&T-mCherry and W-mCherry&T-YFP separately, comparing the resulting parameter estimates with the values used for model simulation (Section S4, Table S2).

### S4 Analysis of microbial traits fitted to competition experiment data

The proposed mathematical model was used to predict the selection dynamics for the competition experiment between a wild-type *E. coli* strain (W) and a tolerant *E. coli* mutant (T) under periodic BAC disinfection. The strains were isolated from a previous laboratory evolution experiment in which the tolerant T evolved from the wild-type W as a result of periodic BAC exposure [1], eventually leading to competitive exclusion of W despite its fitness cost. Here, we used the growth rates of the strains determined in this previous study (without fluorescent tags) to parametrize our mathematical model, while the survival fractions under BAC disinfection were determined through a stationary time-kill assay on the fluorescently labeled strains. Then, for model validation, we assumed that growth rates and disinfection survival during the competition experiment were not significantly affected by fluorescent tags—for cell survival under BAC disinfection, this hypothesis is supported by a two-way repeated-measures ANOVA on the time-kill curves (Section S4). This was assumed to illustrate the use of the approach in practical applications, and to avoid duplicating results (selection and extinction planes, parameter estimates, etc.) for the two competition cases (W-YFP&T-mCherry and W-mCherry&T-YFP), since our objective was to validate experimentally the predictive capabilities of the proposed approach, and not to demonstrate that growth and disinfection survival were independent of fluorescent tags. In other words, to predict selection dynamics using the proposed framework, we only need to isolate the strains from the setting to be disinfected, determine their growth rates and survival fractions, and then derive model predictions, so that fluorescent labeling is not involved in the process.

We performed, nevertheless, an additional model validation to compare the phenotypic traits determined from the previous, independent experiments with those obtained by fitting the model to the competition experiment data for the two competition cases separately. Maximum Likelihood Estimation was used to fit the proposed mathematical model against the competition experiment (flow cytometry) data, assuming independent and identically distributed Gaussian measurements with homoscedastic noise (i.e., constant variance across measurements). That is, the parameter estimates were obtained as those maximizing the Maximum Likelihood Function, which, for independent and identically distributed Gaussian measurements with constant variance, is equivalent to minimizing the unweighted least-squares function between model predictions and measurements [3]. The model parameters 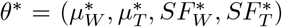 estimated from the competition experiment data were then obtained as:

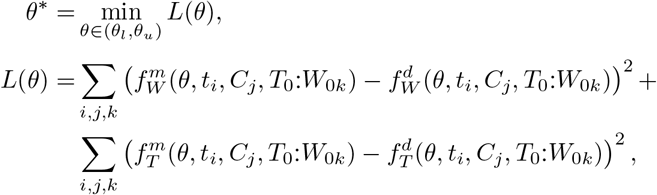

where 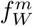 and 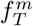 are the subpopulation frequencies predicted by the mathematical model for strains W and T, respectively, and 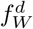 and 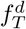 are the subpopulation frequencies obtained from flow cytometry during the competition experiment at sampling times {*t*_*i*_ |*i* = 1, …, *n*_*t*_ } (*n*_*t*_ = 5), BAC doses {*C*_*j*_ | *j* = 1, …, *n*_*C*_ } (*n*_*C*_ = 4), and initial mixing ratios {*T*_0_:*W*_0*k*_ | *k* = 1, …, *n*_*r*_ } (*n*_*r*_ = 3). The lower (*θ*_*l*_) and upper (*θ*_*u*_) bounds on the model parameters used for the optimization problem were set to the confidence bounds for the model parameters in the main text, to provide natural constraints for optimization and further validate if the model satisfactorily fits flow cytometry data within the expected range of variation for the phenotypic traits, determined from previous, independent experiments. The subpopulation frequencies 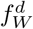 and 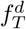 were obtained as the mean of *n* = 3 biological replicates for each competition case (W-YFP&T-mCherry and W-mCherry&T-YFP) separately. Note that the above optimization problem was solved twice, once per competition case.

The model fit shows good agreement with the flow cytometry data (Figure S2 and Table S2). The growth rate of the wild-type W estimated from the competition experiment is slightly lower than that obtained in [1] (0.2 1*/*h) in both competition cases: the estimated value is 1.16 1*/*h when W is labeled with YFP, meaning a reduction of 3% in the value reported before, and the estimated growth rate is 1.12 1*/*h when W is labeled with mCherry, which is around 7% lower than the value reported for the strains without the fluorescent tag. Conversely, the growth rate of the tolerant T estimated from the competition experiment is slightly higher than reported in [1] for the strain without a fluorescent tag (0.96 1*/*h), ranging from 0.98 1*/*h when the strain is labeled with mCherry (2% increase in the value previously reported) to 1.04 1*/*h when it is labeled with YFP (8% increase relative to the value previously reported). These results suggest that fluorescent labeling has a minor impact on growth rates, though the strains labeled with YFP apparently show a slight growth advantage compared to those labeled with mCherry. The value estimated for the fitness cost of the tolerant T thus varies in the range 11 − 17%, compared to a 20% fitness cost for the tolerance mutation reported in [1].

On the other hand, the survival fractions estimated from the competition experiment data are fairly similar to those obtained from the preliminary time-kill assay after joining the data for the two fluorescent labels, though the survival fraction estimated for the tolerant T shows a slightly increase when it is labeled with YFP, except at 30 µg*/*mL BAC. The maximum observed difference is for the survival fraction of the wild-type W labeled with mCherry at 40 µg*/*mL BAC, showing a 13% of reduction (in log10-scale) with the value previously estimated by joining the time-kill curves of the strains for the two labels. The reduced variability observed between the survival fractions estimated for the two competition cases and those obtained from the preliminary time-kill curves is thus attributed to the expected biological variability (as suggested by the two-way repeated-measures ANOVA on the time-kill curves).

Importantly, the variation observed between the phenotypic traits estimated for the two competition cases does not imply a significant variation in the selection dynamics quantitatively predicted by the model for the competition experiment, nor in the predicted experiment outcome (i.e., which strain is eventually selected).

**Figure S2:**
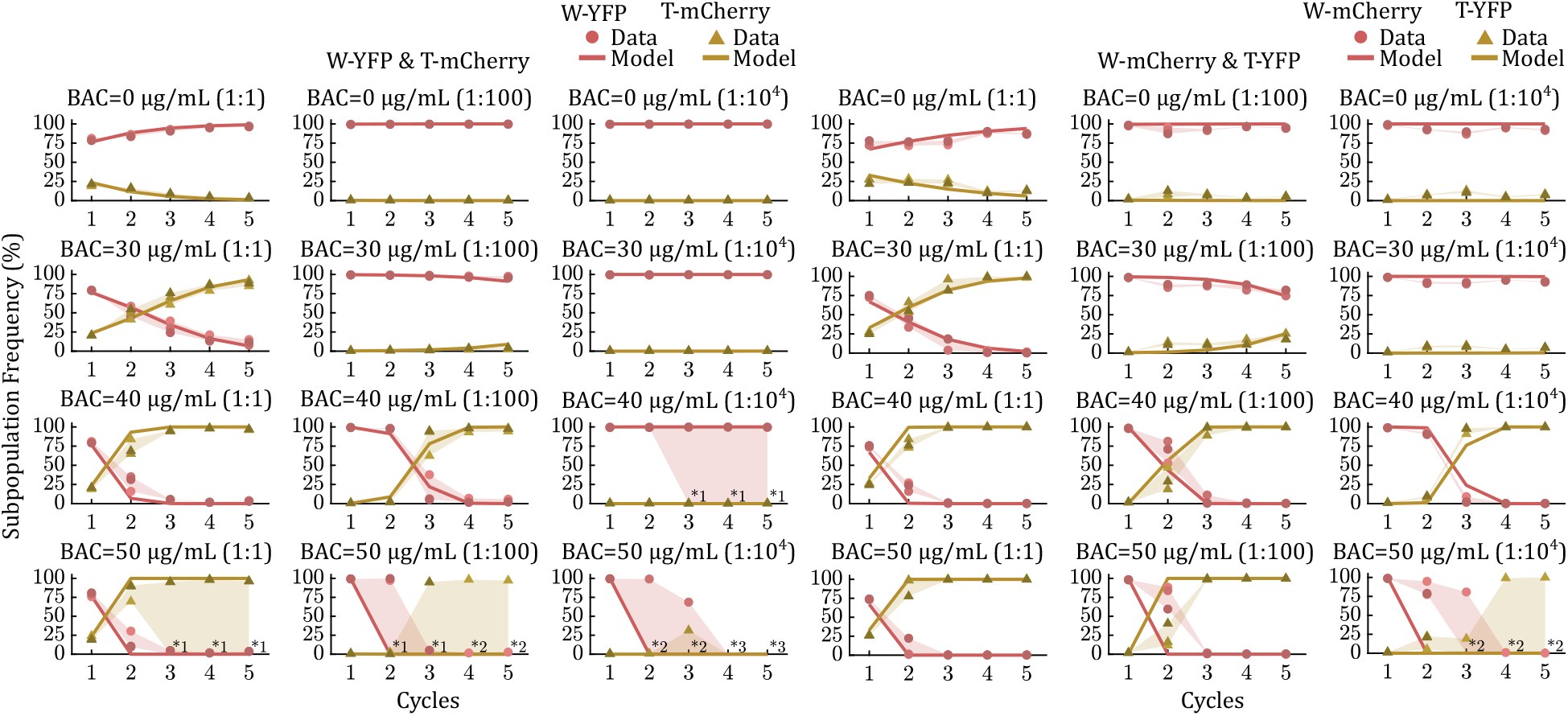
Results of fitting the model for periodic disinfection (see Methods and Table 1 in the main text) to the flow cytometry data (subpopulation frequencies of strains W and T before BAC disinfection) for the competition cases W-YFP&T-mCherry and W-mCherry& T-YFP separately. Data is represented by symbols (dots for strain W and triangles for strain T), and continuous lines represent the model fit. Shading gives the variability between replicates. Total population extinction is indicated by asterisks together with the number of extinct replicates.

**Table S2:** Phenotypic traits (growth rates and survival fractions after 10 min of BAC exposure) of the wild-type (W) and tolerant (T) *E. coli* strains estimated from the competition experiment data. The proposed mathematical model was fitted to the flow cytometry data registered during the competition experiment for cases W-YFP&T-mCherry and W-mCherry&T-YFP separately. The phenotypic traits used to parametrize the model, assuming independence on the fluorescent tag, are also shown for comparison (case Joint).

| Parameter | Symbol | Case |  |  |  |  |  |  |  |  | Unit |
| --- | --- | --- | --- | --- | --- | --- | --- | --- | --- | --- | --- |
|  |  | W-YFP & T-mCherry |  |  | W-mCherry & T-YFP |  |  | Joint |  |  |  |
| Growth rate of W | $\mu_W$ | 1.16 | | | 1.12 | | | 1.20 | | | 1/h |
| Growth rate of T | $\mu_T$ | 0.98 | | | 1.04 | | | 0.96 | | | 1/h |
|  |  | BAC |  |  |  |  |  |  |  |  |  |
| | | 30 | 40 | 50 | 30 | 40 | 50 | 30 | 40 | 50 | $\mu\text{g/mL}$ |
| Survival fraction of W | $\log_{10}(SF_W)$ | -1.11 | -3.55 | -7.21 | -1.01 | -3.91 | -7.20 | -1.13 | -3.45 | -7.24 | — |
| Survival fraction of T | $\log_{10}(SF_T)$ | -0.31 | -1.66 | -4.64 | -0.36 | -1.38 | -3.99 | -0.27 | -1.66 | -4.35 | — |
| Root mean square error | RMSE | 0.28 |  |  | 0.22 |  |  | — |  |  |  |

### S5 Detection of total extinction during the competition experiment

The competition experiment performed in this work consisted of mixing the wild-type *E. coli* strain (W) and the tolerant *E. coli* mutant (T), and monitoring the selection dynamics between the two strains over successive growth and disinfection periods with benzalkonium chloride (BAC). Four complete cycles of growth and disinfection were performed, followed by an additional growth period, concluding the experiment. The subpopulation frequencies of the strains within the mixed cultures were monitored with flow cytometry before each disinfection round. Additionally, we plated serial dilutions of the mixed cultures to determine the concentration of viable cells before each disinfection round (Figure S3). We then combined the cell count data and the flow cytometry data to assess extinction of the replicates for the competition experiment; extinction of a biological replicate were thus considered when no viable cell counts were detected above the low detection limit before disinfection starts, and these results were compared with the number of events detected in flow cytometry (Figure S4). The extinction of the mixed cultures should be assessed before disinfection, as there is no guarantee that the population will not regrow in the subsequent growth period when the cell density falls below the low detection limit after disinfection. After the last growth period, additional measurements were collected using flow cytometry, but not serial plating.

**Figure S3:**
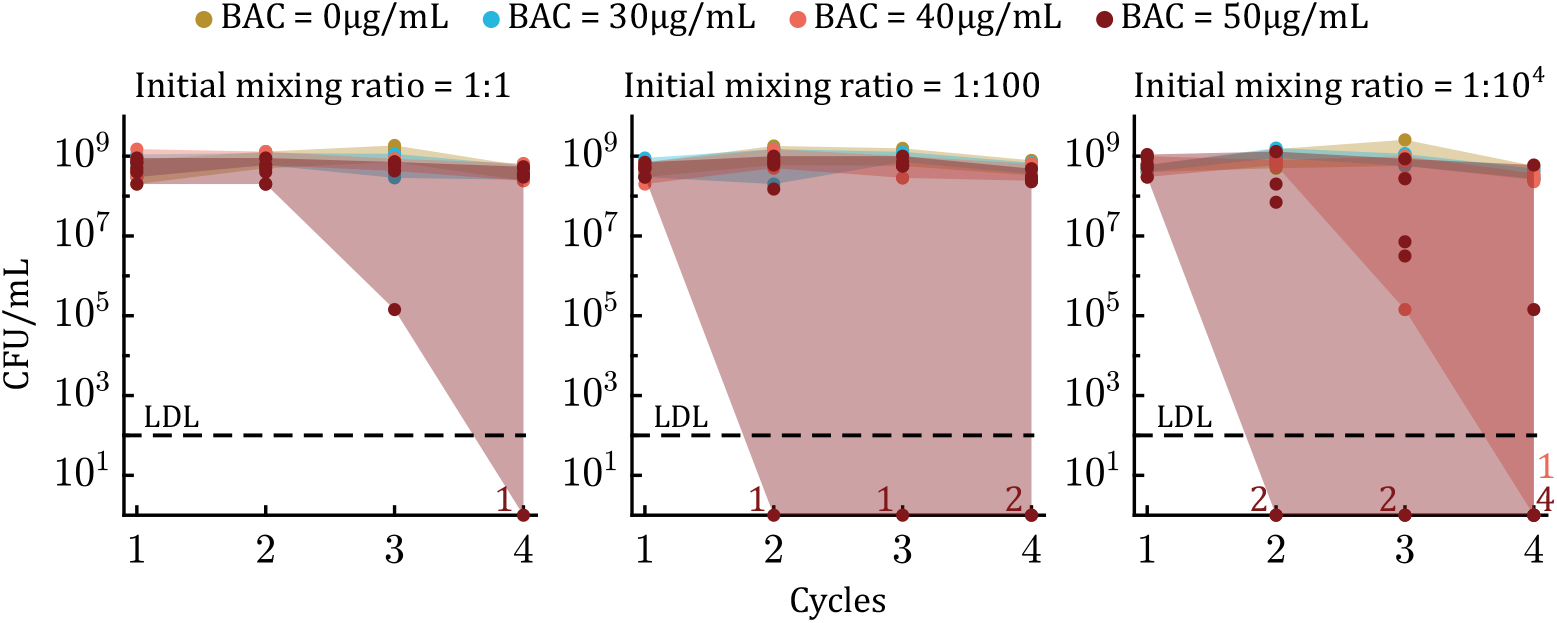
Total cell counts measured during the competition experiment between the wild-type *E. coli* strain (W) and the tolerant *E. coli* mutant strain (T) at the start of each disinfection round, determined by plating serial dilutions of the mixed cultures. Data is represented by dots for the distinct BAC concentrations and initial mixing ratios tested in the competition experiment, and shading gives the variability range between biological replicates. Strains W and T were fluorescently labeled with YFP or mCherry to measure the subpopulation frequencies with flow cytometry during the competition experiment, obtaining two competition cases (W-YFP&T-mCherry and W-mCherry&T-YFP), with *n* = 3 biological replicates each (per BAC dose and initial mixing ratio). Data from the two competition cases were combined and shown together, resulting in *n* = 6 biological replicates. Asterisks indicate extinction of the replicates, i.e., total cell density below the low detection limit (LDL = 100 CFU*/*mL), together with the number of extinct replicates.

The results obtained from flow cytometry and cell counting were fairly consistent, so that the replicates for the competition experiment that were classified as extinct using the criterion based on cell count data coincide with these replicates showing a drastic decrease in the number of events detected by flow cytometry (compared with the control case in which the mixed cultures are not treated with BAC). There were only a few discrepancies between the cell count data and the flow cytometry data. First, at the highest initial mixing ratio (*T*_0_:*W*_0_ = 1:1), the unique extinct replicate at 50 µg*/*mL BAC was detected one cycle before by flow cytometry than by cell counting. Second, at the lowest initial ratio (*T*_0_:*W*_0_ = 1:10^4^), flow cytometry detected extinction of one of the replicates one cycle before cell counting at 40 µg*/*mL BAC; at 50 µg*/*mL BAC, similarly, there are also two replicates that were classified as extinct by flow cytometry one cycle before than by cell counting. In these cases, we used the results from flow cytometry to establish extinction of the replicates, since this technique is more reliable and less prone to experimental error than cell counting.

It should be noted that population extinction during periodic disinfection was determined from the mathematical model and from the data in different (though comparable) ways, as the concept of population extinction (or disinfection success) vary depending on the application and on the experimental techniques. The proposed mathematical model considers that a strain is extinct when its cell density falls below an extinction limit—set to *X*_*e*_ = 1 CFU*/V* in this work, where *V* is the culture volume—after disinfection periods. From this point, the strain is not allowed to reproduce during the following growth periods, so that its cell density remains below the extinction limit for the remainder of the model simulation. That is an advantage of mathematical models, as they allow population dynamics to be captured at lower densities than can be detected by, e.g., plating serial dilutions. On the other hand, flow cytometry allows analysis of cell properties individually and works with small populations, but it is not straightforward to assess cell viability with this technique. For the mathematical model, we assumed an extinction threshold determined by biological constraints (the population cannot reproduce if there are fewer than 1 CFU in the culture volume *V*) rather than by experimental constraints. Nevertheless, the distinct extinction thresholds used in this work, both for model and data, yielded consistent results, so that neither of the data replicates classified as extinct showed regrowth after falling below the extinction threshold for cell counting or flow cytometry. Furthermore, the model satisfactorily predicted total population extinction when more experimental replicates were classified as extinct by the experimental criterion.

### S6 Dependence of the extinction plane on the fitness cost

The mathematical modeling framework proposed in this work was used to derive extinction conditions for assessing the success of periodic disinfection (see Methods in the main text). These extinction conditions involved four distinct phenotypic raits: growth rates in the absence of disinfection and survival fractions after disinfectant exposure for the two competing strains, wild-type (W) and tolerant (T). Then, the extinction conditions cannot be represented against the phenotypic traits for the strains separately, because extinction is influenced by a competitive part linking the dynamics of the two competing strains. That is, a strains that would survive periodic disinfection in isolation (i.e., independently of competition with the other) can be eliminated in just a single treatment cycle in competition. Thus, we could represent an extinction plane for the strains in isolation as a function of growth rate and survival fraction, but we would miss the competitive part of the extinction dynamics. Also, the extinction conditions do not depend solely on the fitness cost and survival advantage of the tolerant T, which are relative measures comparing selection between the strains, but rather on the phenotypic traits of strains W and T separately.

**Figure S4:**
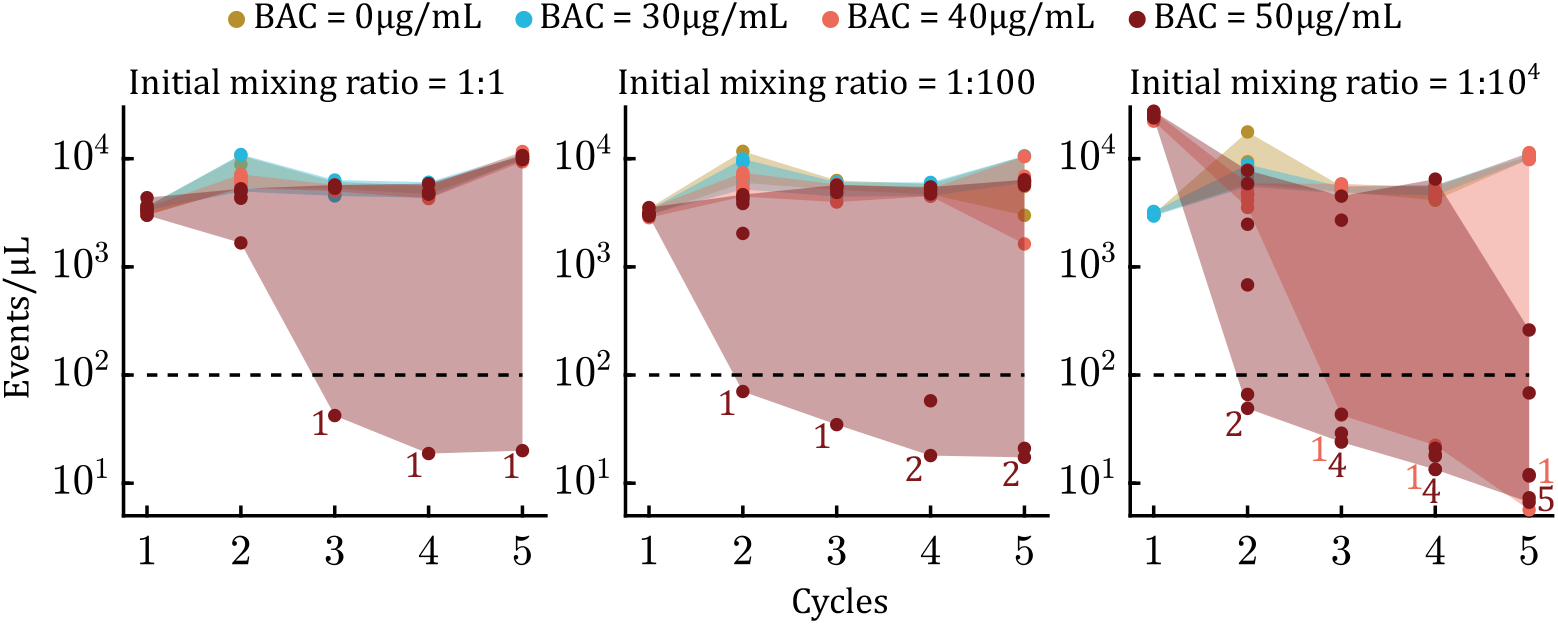
Number of events per micro-liter detected with flow cytometry during the competition experiment between the wild-type *E. coli* strain (W) and the tolerant *E. coli* mutant strain (T) at the start of each disinfection round. Data is represented by dots for the different BAC doses and initial mixing ratios tested in the competition experiment, and shading gives the variability range between biological replicates. Strains W and T were fluorescently labeled with YFP or mCherry to measure the subpopulation frequencies with flow cytometry during the competition experiment, obtaining two competition cases (W-YFP&T-mCherry and W-mCherry&T-YFP) with *n* = 3 biological replicates each (per BAC dose and initial mixing ratio). Data from the two competition cases were combined and shown together, resulting in *n* = 6 biological replicates. Asterisks represent biological replicates for which flow cytometry detected less than 100 events per micro-liter, together with the number of such replicates. The figure only considers detected events classified as bacteria from one of the competing strains (W and T) after preprocessing and clustering the raw flow cytometry data.

Therefore, the extinction conditions cannot be fully represented in a plane, as done with the selection conditions, because they involve more than two microbial parameters whose influence on the extinction dynamics cannot be decoupled. Since the extinction conditions for wild-type W and tolerant T competing under periodic disinfection cannot be represented on a plane, we worked in the main text with a reduced extinction plane where the growth rates of the strains were set to the values estimated from previous, independent experiments. This procedure enabled the representation of model-based predictions of strains’ extinction at different disinfectant doses while maintaining a constant value for the fitness cost. Nevertheless, we are not limited to analyzing the extinction scenarios when assuming the estimated fitness cost for the tolerant T; we can also assess extinction in scenarios where the tolerant T shows the minimum and maximum fitness cost allowed by the confidence bounds on the growth rates (Figure S5). As a result, we would obtain a range of extinction scenarios for the periodic disinfection protocol, allowing us to analyze what would happen when the tolerant T evolves as the minimum fitness cost allowed, for example.

Figure S5 illustrates the dependence of the extinction conditions on growth rates, and thus on the fitness cost of the tolerant T. The bounds for extinction of strains W and T in competition are not affected by the variation in growth rates, because the duration of the growth periods for the competition experiment is long (*t*_*g*_ = 24 h). That is, the strains in isolation reach the carrying capacity (*K*) in the first cycle even when they grow at the minimum allowed growth rate, so that the extinction bounds are entirely determined by the bottleneck density set by carrying capacity and dilution (i.e., *DK*). Conversely, the extinction bounds for the strains in competition show a strong dependence on growth rates (and on strains’ inocula), as these traits determine selection during periodic disinfection when strains W and T compete for growth-limiting resources. That is, a strain can survive periodic disinfection in isolation but go extinct during competition if it is at a competitive disadvantage (determined by growth rates and inocula) during the first growth period, so it cannot consume enough resources to survive after disinfection and dilution, and thus propagate into the next cycle.

**Figure S5:**
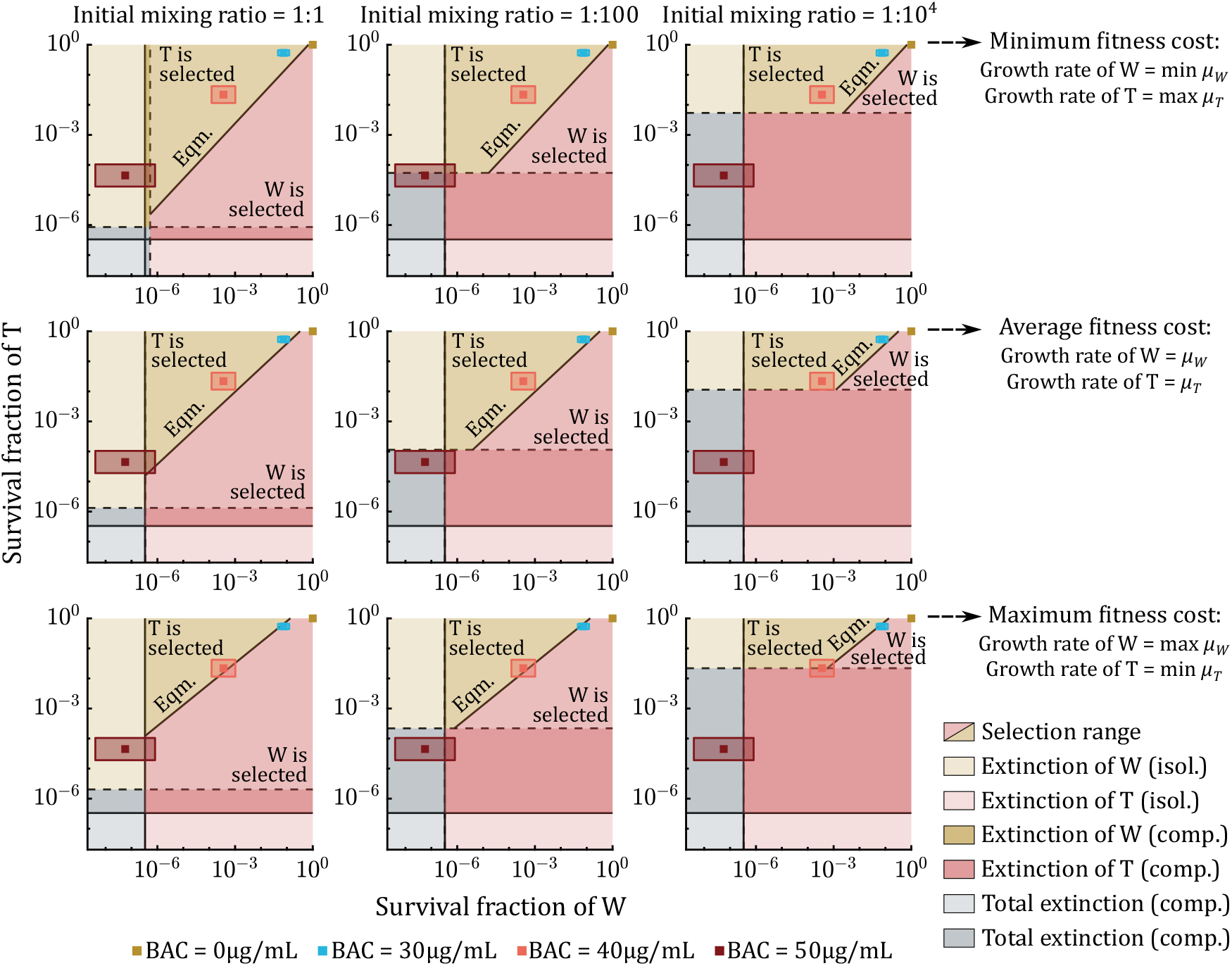
Reduced extinction plane for the competition experiment between the wild-type (W) and the tolerant (T) *E. coli* strains under periodic BAC disinfection, considering the confidence bounds for the growth rates of the strains, and thus a confidence range in the fitness cost of T. The extinction plane is represented by assuming the minimum fitness cost (top) obtained from the confidence bounds on the growth rates, compared with the estimated (middle) and maximum (bottom) values. The different regions on the extinction plane indicated in the legend are determined by lower bounds on the survival fractions of strains W and T, both in isolation (continuous lines) and in competition (dashed lines). Squares show the position on the extinction plane predicted from the extinction conditions with the estimated survival fractions of the strains at the different BAC doses, and the shaded boxes represent the confidence bounds on the estimates.

The results in Figure S5 show that the fitness cost of the tolerant T significantly influences the outcome of periodic disinfection. When the fitness cost is assumed to be maximum, we observe scenarios on the reduced extinction plane where the wild-type W is selected over the tolerant T and survives periodic disinfection; the strains are close to the selective equilibrium at 40 µg*/*mL BAC, while strain W outcompetes T for lower BAC concentrations. This is especially noticeable when wild-type W and tolerant T start periodic disinfection from equal competition (*T*_0_:*W*_0_ = 1:1). In this case, the wild-type W exerts the strongest competence on the tolerant T during the first cycles of treatment, so that we can observe scenarios where strain T is outcompeted by W at all tested BAC doses when we assume the maximum fitness cost of tolerance. Conversely, when the fitness cost of T is set at its minimum, the tolerant T is preferably selected over the wild-type W within the uncertainty range for the survival fractions at all tested BAC concentrations.

Figure S5 can also be used to analyze the uncertainty in the prediction of total population extinction obtained at the highest BAC dose (50 µg*/*mL) from the estimated fitness cost of strain T. Indeed, the extinction conditions assuming the estimated value of fitness cost predicted total population extinction for *T*_0_:*W*_0_ = 1:100, 1:10^4^ (i.e., when there are not enough T cells initially present in the population). However, for *T*_0_:*W*_0_ = 1:100, we can now observe that population survival is feasible within the uncertainty range of the survival fractions in the worst-case scenario where the fitness cost of the tolerant T is assumed to be the minimum. We can therefore consider this uncertainty in the extinction conditions arising from the fitness cost of the tolerant T to yield worst-case predictions for which there may be some probability of disinfection failure. On the other hand, note that the model predictions of total population extinction from the estimated fitness cost are maintained at *T*_0_:*W*_0_ = 1:10^4^, indicating that total population extinction is almost sure within the uncertainty range of the fitness cost in this case.

## Notes

https://zenodo.org/records/19910150

### Competing Interest Statement

The authors have declared no competing interest.

